# A Toolkit for Rapid Modular Construction of Biological Circuits in Mammalian Cells

**DOI:** 10.1101/506188

**Authors:** João Pedro Fonseca, Alain R. Bonny, G. Renuka Kumar, Andrew H. Ng, Jason Town, Qiu Chang Wu, Elham Aslankoohi, Susan Y. Chen, Patrick Harrigan, Lindsey C. Osimiri, Amy L. Kistler, Hana El-Samad

## Abstract

The ability to rapidly assemble and prototype cellular circuits is vital for biological research and its applications in biotechnology and medicine. Current methods that permit the assembly of DNA circuits in mammalian cells are laborious, slow, expensive and mostly not permissive of rapid prototyping of constructs. Here we present the Mammalian ToolKit (MTK), a Golden Gate-based cloning toolkit for fast, reproducible and versatile assembly of large DNA vectors and their implementation in mammalian models. The MTK consists of a curated library of characterized, modular parts that can be easily mixed and matched to combinatorially assemble one transcriptional unit with different characteristics, or a hierarchy of transcriptional units weaved into complex circuits. MTK renders many cell engineering operations facile, as showcased by our ability to use the toolkit to generate single-integration landing pads, to create and deliver libraries of protein variants and sgRNAs, and to iterate through Cas9-based prototype circuits. As a biological proof of concept, we used the MTK to successfully design and rapidly construct in mammalian cells a challenging multicistronic circuit encoding the Ebola virus (EBOV) replication complex. This construct provides a non-infectious biosafety level 2 (BSL2) cellular assay for exploring the transcription and replication steps of the EBOV viral life cycle in its host. Its construction also demonstrates how the MTK can enable important and time sensitive applications such as the rapid testing of pharmacological inhibitors of emerging BSL4 viruses that pose a major threat to human health.

## Introduction

Molecular cloning is the cornerstone of modern biological research. It is required to generate DNA vectors that can encode a myriad of genetic circuits with applications in the fields of cell biology, therapeutics, virology, synthetic biology and biotechnology, among others. These circuits can encompass the expression of endogenous proteins, small RNAs, reporters of gene expression or cellular activity, and other innumerable reagents whose applications extend from basic discovery to medical applications. To move molecular cloning forward into an engineering discipline, with minimal trial and error and reproducible results, the parts that constitute engineered circuits need to be modular, vetted for their function and easy to share. To maximize the range of applications, circuits made from these parts need to be rapidly assembled, amenable to fast prototyping, and not bottlenecked by the delivery method to their target genomic integration sites.

Conventional approaches to produce DNA vectors involve the PCR amplification of DNA fragments from different sources, which are later ligated following digestion with restriction enzymes. This procedure is time consuming, laborious, prone to errors and requires many control steps when subcloning (e.g. sequencing). Due to the shrinking cost of commercial DNA synthesis, ordering gene fragments or entire genes provides a potential avenue to reliably build libraries of DNA vectors. While this technology increases the diversity of parts that can be cloned and removes a DNA source requirement, current pricing, size limitation, and turnaround time remain a significant bottleneck.

In light of these limitations, several strategies have been developed to streamline the assembly of different parts in the construction of DNA vectors for mammalian biology^1–7^. Notable among those is the Gibson modular assembly platform (GMAP)^7^, a framework for DNA vector construction by Gibson assembly. More recently, the Mammalian Modular Cloning (mMoClo)^6^ and Extensible Mammalian Modular Assembly kit (EMMA)^3^ based on Golden Gate assembly reactions were proposed. Both methods use sets of modular parts with given functionality, which can be assembled directionally into transcription units (TUs) by type-IIS restriction enzymes. While useful, all three methods have important shortcomings. GMAP requires sequencing in between all cloning steps, which makes the method slow and expensive. EMMA and mMoClo provide a method to hierarchically assemble DNA vectors that encode large circuits, but fall short on shareability, integration of Cas9 sgRNA cloning, a library of tested parts, and ease of use. While in yeast and plant systems, methods for the rapid, efficient and hierarchical assembly of genetic circuits from a large library of well described modular parts^8–11^ exist, we are unaware that such a cloning toolkit for mammalian systems is readily available.

In this work, we adapted the modular cloning strategy (MoClo)^12^ strategy to build mammalian expression vectors. The Mammalian ToolKit (MTK) is a library of over 300 parts including vetted promoters, 3’UTRs, fluorescent proteins, insulator and P2A elements that can be combined to build large genetic circuits in a rapid and efficient fashion. These vectors can be delivered to a wide range of cell types, through viral, recombinase and CRISPR/Cas9 methods. As a proof of concept, we built a hAAVS1 landing pad for single integration of genetic circuits, created and delivered libraries of protein variants and sgRNAs and rapidly compared multi-species Cas9-based genetic circuits. Finally, we used Ebola virus (EBOV) as a test case to address an emerging need to streamline the generation of BSL2 reagents during outbreaks of novel BSL4 viruses. Using the MTK, we rapidly built a non-infectious multicistronic ribonucleoprotein expression system for EBOV in a mammalian host. This system recapitulates the transcription and replication sub-lifecycle of the virus in a mammalian host, and can therefore be used to accelerate the study of viral biology and discovery of antiviral molecules.

## Results

### An expansive, modular cloning toolkit for rapid prototyping in mammalian cells

The basis for the MTK is a library of parts that are “domesticated” from source DNA using a Golden Gate (GG) reaction^12^ into a standard vector. The resulting plasmid is then sequenced once to ensure fidelity to the source sequence, and then can later be reused as a modular part in a variety of genetic constructs. Similar to previously published work ^6,3,10,12^, we utilize the type-IIS restriction enzymes BsaI and BsmbI due to their “reach over” endonuclease activity that leaves arbitrary four base overhangs adjacent to the recognition site. The defining feature for a part vector is a unique four base overhang that categorizes it to similar parts, and ensures the sequential 5’ to 3’ ligation of parts into a transcriptional unit (TU) of multiple parts. Due to the hierarchical nature of the MTK, with the library of part vectors sequenced-verified, only diagnostic restriction digests are necessary to verify correct assembly in subsequent GG reactions. We employ BsmBI and BsaI GG reactions in an alternating manner, to assemble a library of part vectors into libraries of TU plasmids, and ultimately into multi-transcriptional unit (multi-TU) plasmids. The 4 day time from source DNA to final plasmid is comparable to alternative cloning methods when including time to sequence verify (Fig. 1a). However, the significant time reduction comes in the reusability of the confirmed part plasmids where completely redesigned new configurations of final plasmids (e.g. new TUs) can then be assembled from existing parts in only 2 days.

**Fig. 1.**
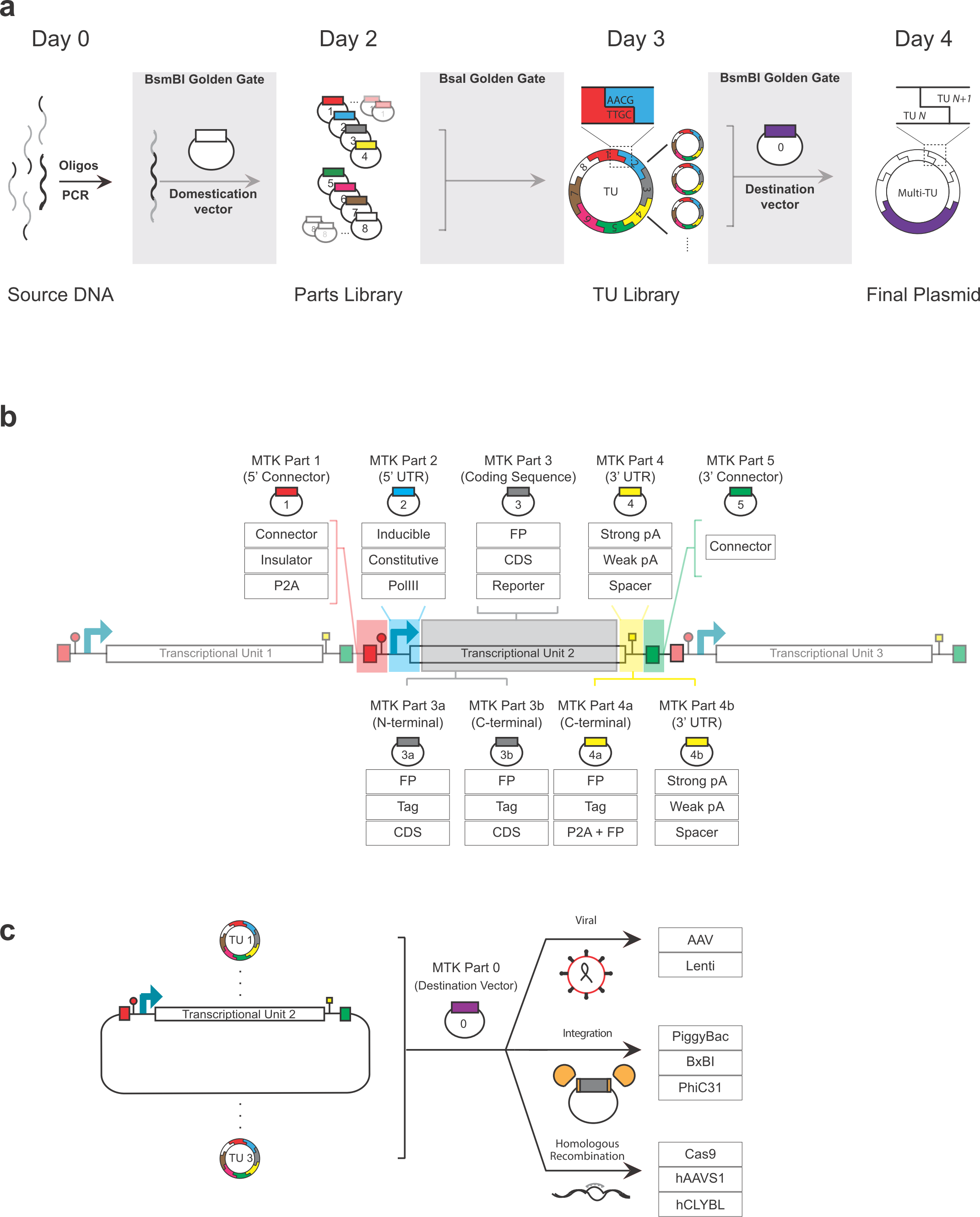
Schematic and definition of parts of the Mammalian Toolkit (MTK). **(a)** Workflow of the MTK starting with a BsmBI part domestication, BsaI transcriptional unit assembly, and a final BsmBI assembly of a multi-transcriptional unit plasmid. Top text indicates approximate time from initial PCR with primers to receiving final plasmid (**b**) Schematic of a standard transcriptional unit (TU). Tables below part definition summarize variations of that part that are present in the MTK. Supplemental table 1 contains a comprehensive list of current parts (**c**) Library of TUs can be re-used with no further modification and delivered to cells in three methods already present in the MTK using a dedicated destination vector (MTK part 0).

The MTK encompasses part vector categories 1-8 that are sufficient to build and deliver a vast combinatorial library of genetic constructs to cells (Fig. 1b). Parts 2, 3 and 4 form the core of the TU, specifying the 5’ UTR, coding, and 3’ UTR sequences, respectively. Part 2 corresponds to promoter sequences that can be specified to drive variable constitutive or inducible expression or recruit diverse host polymerases, such as polymerase III for non-coding RNA transcription. Part 3 vectors are canonical coding sequences that are typically proteins of interest. Part 4 refers to 3’ UTR sequences that encode polyadenylation (pA) sequences that terminate transcription, or spacer sequences that couple transcription to the downstream TU. Finally, parts 1 and 5 correspond to connector sequences that enable the sequential ordering of TUs into a multi-TU plasmid, with many versions provided in the MTK. For example, if the design objective is to minimize polymerase read-through and maximize independent expression from each TU, then Part 1 implements connectors with insulator sequences^13^. Conversely, if the goal is to enable strongly correlated expression across adjacent TUs, alternative Part 1 vectors consisting of connectors with orthogonal P2A elements are provided. These are cis-acting hydrolase elements that allow the promoter of one TU to drive expression of up to five downstream TUs^14^.

Within these defined parts, further specialization is easily achieved. For example, Part 3 and Part 4 vectors can be replaced with constituent ***a*** and ***b*** vectors where Part 3 can be replaced with Part 3a and 3b, and still connect to a Part 4. This allows combinations that can implement tethering of localization tags, protein domains or any desired coding sequence both N-and C-terminally with an innocuous linker sequence in between (Fig. 1b). Additionally, we have developed coupled Part 234 vectors to accommodate rapid cloning of small guide RNA (sgRNA) expression by oligo annealing for CRISPR/Cas9-related genetic constructs. Parts 6, 7 and 8 generally flank a typical TU and can encode the method of delivery to cells. For example, it can encode the homology arms for a locus of integration, or bacterial origin of replication and selection markers.

Lastly, the MTK contains Part 0 destination vectors that allow delivery via viral transduction, transposase transfection and homologous recombination from the same collection of TUs (Fig. 1c). Unlike other assembly methods, where the genetic constructs must be repeatedly redesigned to accommodate a particular delivery method, within the MTK, one needs only to change the Part 0 vector in the final GG reaction to facilitate different delivery into cells. Several considerations may determine which destination delivery is appropriate, but this system allows one to recycle TUs and maintain a library of useful units that may be context-specific. This mammalian cloning toolkit combines intuitive, hierarchical organization, with an expansive library to enable rapid, facile iteration between a variety of genetic constructs that can be integrated into cells without redesign or re-sequencing, irrespective of the delivery method. A comprehensive list of parts can be found in Supp. Table 1.

### MTK enables facile construction of TUs with different levels of expression

In designing genetic constructs, an important consideration is the relative doses of expression strengths between respective genes. Within the MTK, we have seventeen characterized constitutive promoters that span a spectrum of transcriptional expression strengths. We include promoters from a mix of human, mouse and viral origin for a range of expression, alternatives to swap in the event of silencing effects, and a balance of native and transgene promoters for conventional cell lines.

To characterize the relative strengths of these promoters, we assembled a panel of 14 TUs with varied promoter parts driving mAzamiGreen expression, flanked by the bovine growth hormone (Bgh) pA 3’ UTR. In an adjacent TU, a CAG promoter expresses mScarlet with the rabbit beta-globin (Rgl) pA 3’ UTR (Fig. 2a). We transiently transfected each multi-TU plasmid into HEK293T cells and normalized the mAzamiGreen expression by mScarlet to control for plasmid copy number. This suite of promoters span a range of 2 orders of magnitude over background, with the strongest promoter, CMV, more than 300-fold greater than a promoter-less mAzamiGreen. The ranked expression strength of the promoters revealed a smooth continuum of expression between 10-fold and 300-fold (Fig. 2a). Thus this suite of promoters can be used to tailor TU expression levels and leverage overlapping promoter strengths to avoid repeated elements in multi-TU plasmids. The ranked expression of the promoters across human and mouse demonstrates that their relative strengths are largely maintained (Fig. Supplementary Fig. 1a, b), suggesting that these promoters are portable across different systems. Additionally, the rank of promoter effect on gene expression is consistent between transient and stably integrated expression, leaving this suite of promoters as a reliable metric for TU design (Fig. Supplementary Fig. 1c, d).

**Fig. 2.**
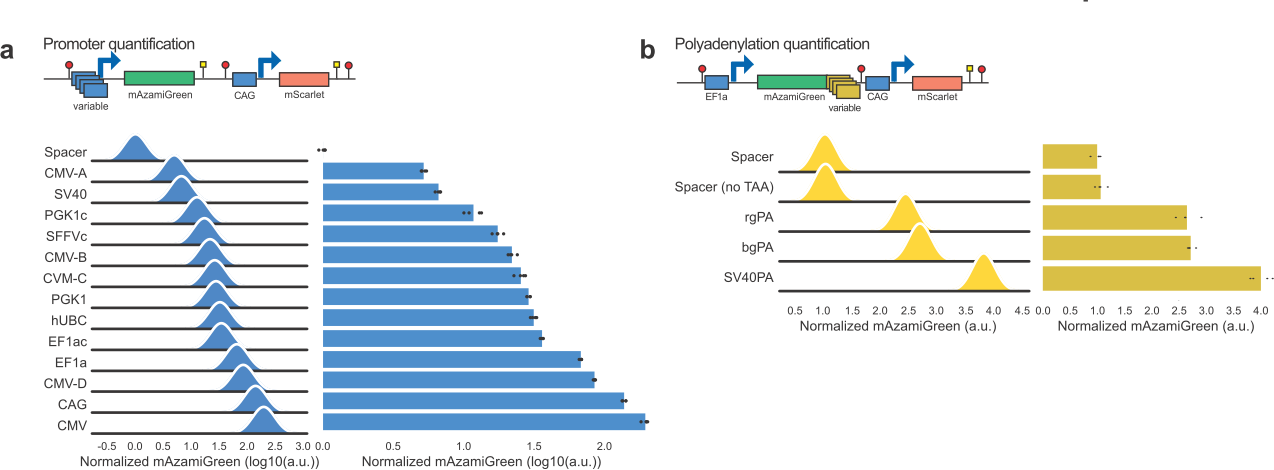
Characterization of constitutive promoters and 3’ UTR provided in the MTK. **(a)** Top: Schematic of a TU used to quantify promoter expression. Bottom: Ranked expression of mAzamiGreen (normalized to a constitutive mScarlet) driven by different promoters. Distributions show one of four biological replicates, and bar plots represent the mean of all four biological replicates. The mean of each replicate is also shown as a black dot. The letter “c” following the name of a promoter (e.g PGK1 versus PGK1c) is used to designate a “crippled” promoter where the Kozak sequence is disrupted. (**b**) Top: Schematic of TU used to quantify the effect of the 3’ UTR on mAzamiGreen expression driven by the EF1a promoter. Data representation is as described above.

Additionally, the MTK allows for further fine-tuning of a TU in a given range using a collection of five different 3’ UTR sequences. Similar to the promoter comparison, we generated a circuit with the same constitutive promoter (EF1a) and varied the 3’ UTR sequence to compare normalized mAzamiGreen expression (Fig. 2b). We observed a range of 4-fold change in expression of mAzamiGreen among the three conventional 3’ UTR sequences. While the Bgh pA and Rgl pA signals are redundant, the simian virus 40 (SV40) pA signal has a nearly 1.5-fold greater effect on expression than the other pA sequences (Fig. 2c). This supports the notion that by rationally varying promoter-pA combinations, one can fine-tune incremental changes in expression to complement the log-scale changes accompanied by solely changing promoters. Moreover, replacing a canonical 3’ UTR sequence with either of the spacer sequences diminishes expression of its upstream coding sequence, but the spacers still enable appreciable expression over background. The two spacers differ in their use: including a stop codon allows for efficient reverse-transcription in lentiviral preparation while omitting the stop codon enables read-through in multicistronic plasmids. Finally, the MTK contains two inducible promoters whose use is illustrated in Fig. 6. This characterized suite of constitutive promoters and 3’ UTR’s coupled with the facile assembly can generate libraries of fine-tuned expression of a protein of interest. We only explored here a few combinations related to one promoter (EF1a), but other combinations can be easily assembled using the parts already provided in the MTK and assayed by the interested user.

**Fig. 6.**
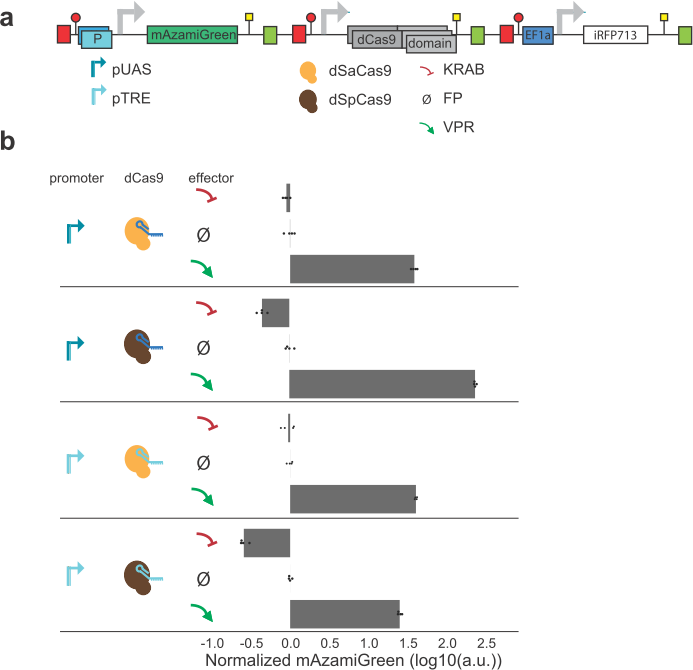
Parallelization of dCas9 circuit prototyping using the MTK. **(a)** Schematic of 12 dCas9 circuit variations assembled in parallel. Each variation is a multi-TU plasmid with either a TRE or UAS inducible promoter controlling the expression of mAzamiGreen, followed by EF1a driving either dSpCas9 or dSaCas9 fused to a KRAB domain, fluorophore, or VPR domain. UAS, Upstream Activating Sequence.TRE, Tetracycline Responsive Element. dSpCas9 and dSaCas9, deactivated S. *pyogenes* and S. *aureus* Cas9, respectively. KRAB, Krueppel-associated box. FP, fluorescent protein. VPR, VP64-p65-Rta. **(b)** mAzamiGreen expression (normalized by iRFP713) for different configurations of the circuit. Every row is a different configuration, corresponding to a variation of the promoter, dCas9 used, and effector used. Configurations are grouped by their promoter-dCas9 pairing (different groups are separated by solid lines). Bar plot represents mean of 4 biological replicates, the mean of each is shown as a black dot.

### MTK contains linkers encoding P2A elements for building multicistronic transcriptional units

One hurdle to construct genetic circuits for viral delivery is that viral vectors have a limited cargo capacity. While this issue can be bypassed by using other delivery methods, if viral delivery is the only option, smaller sized constructs that maintain function are needed. To address this need, we developed Left Connector parts (MTK Part 1) that encode ribosome skipping P2A elements to allow for the expression of 2-12 genes of interest in a multicistronic vector. To test the toolkit’s ability to produce skipped proteins from one RNA, we built a multicistronic vector with one CAG promoter driving membrane-tethered iRFP713, cytoplasmic mAzamiGreen, mCerulean-tagged p38 KTR^15^, and histone H2B fused to mScarlet separated by P2A sites (Fig. 3a). Compared to the control version of this construct, which expresses the same proteins in a multi-TU format from repeated, independent promoters, the multicistronic construct conferred a 35% reduction in vector size. Transiently transfected HEK293T cells with the multicistronic vector showed the correct localization of each of the proteins (Fig. 3b), with no distinguishable difference from the independent multi-TU control. These data demonstrate the feasibility to deploy P2A elements to generate single multicistronic constructs that circumvent the size limitations of conventional viral delivery vectors.

**Fig. 3.**
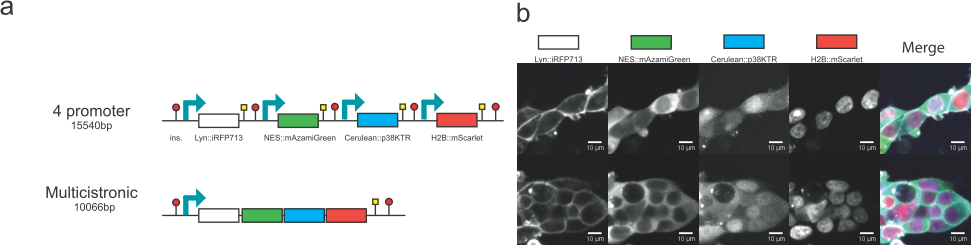
A multicistronic construct allows the expression of 4 differentially located fluorescent proteins. **(a)** Schematic of expression strategies of 4 fluorescent proteins targeted to 4 different cellular locations. In the first strategy, a plasmid contains 4 TUs, each containing a CAG promoter. In the first TU, CAG drives expression of membrane-targeted iRFP713 (Lyn::iRFP713), in the second expression of cytoplasmic mAzamiGreen (NES::mAzamiGreen), in the third expression of p38 kinase translocation reporter fused to mCerulean (Cerulean::p38KTR), and in the fourth histone H2B fused to mScarlet (H2B::mScarlet). In the second strategy, a multicistronic plasmid encodes the same proteins, but all are produced from a single transcript driven by a CAG promoter with 3 P2A sequences separating the 4 peptides. The size in bp of each plasmid is also indicated **(b)** Confocal images of HEK293T cells expressing the 4 promoter (top) or multicistronic (bottom) plasmid. Merge panel shows Lyn::iRFP713 in white, NES::mAzamiGreen in green, Cerulean::p38KTR in blue and H2B::mScarlet in red.

### MTK contains landing pad system and accompanying destination vector

When delivering synthetic genetic circuits, it can be essential to have site-specific, single copy integration. In addition, it is important that this integration is efficient and reliable. In order to accomplish this functionality, we included in the MTK a BxBI-dependent landing pad system for integrating synthetic circuits in a locus of choice. The landing pad system is divided into two parts: first, a landing pad cell line; and second, a landing pad transfer vector. As a proof of concept we built a HEK293T cell line with a BxBI landing site in the hAAVS1 locus (hAAVS1 LP), identified by mRuby2 expression and Hygromycin resistance, and tested its ability for site specific recombination.

The landing pad cell line was generated by CRISPR/Cas9-mediated integration of a vector with homology to the hAAVS1 locus. This vector encoded a multicistronic cassette with Hygromycin resistance and nuclear-localized mRuby2 surrounded by a LoxP site and two FRT sites. The BxBI attP site is located between the promoter and the first gene of this cassette (Fig. 4a). We chose to integrate the landing pad in the hAAVS1 locus due to its low impact on gene expression^16^. We verified the correct integration of the landing pad cassette by PCR in 8 clones (Supplementary Fig. 3) and continued its characterization in clone #8 due to the monoallelic presence of the landing pad. We further confirmed the presence of the landing pad in the cell line by mRuby2 expression (Fig. 4b).

**Fig 4.**
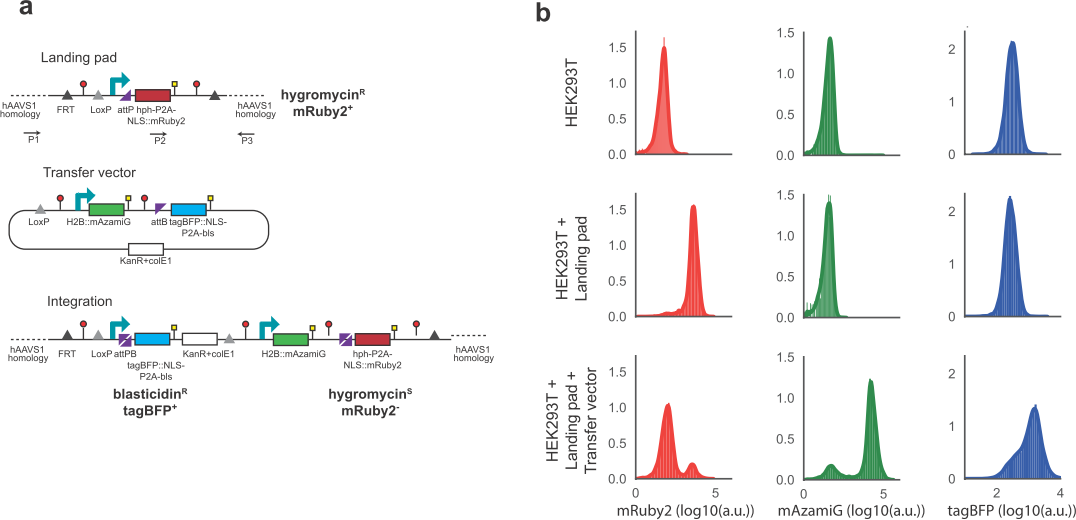
Generation and testing of of landing pads for HEK293T cells using the MTK. **(a)** Schematic of landing pad, transfer vector and integrated vector. Also shown are genotyping primers. FRT, flippase recognition target; ins., Insulator; LoxP; LoxP site; attP, BxBI phage attachment site; hph, Hygromycin resistance gene, bls, Blasticidin resistance gene; NLS, nuclear localization signal; KanR, Kanamycin resistance gene; colE1, colE1 origin of replication; attB, BxBI bacterial attachment site; mAzamiG, mAzamiGreen. **(b)** mRuby2, mAzamiGreen and tagBFP expression in populations of parental, Landing pad and Landing Pad with Transfer vector HEK293T cells. mRuby2 expression indicates presence of hAAVS1 landing pad. mAzamiGreen and tagBFP expression indicates precise integration of transfer vector in hAAVS1 landing pad.

The second part of the landing pad system is a transfer vector that encodes the synthetic gene circuit. In order to verify that the transfer vector is correctly integrated at the hAAVS1 landing pad (hAAVS1 LP), we positioned a promoter-less multicistronic cassette encoding Blasticidin resistance and nuclear localized tagBFP downstream of the BxBI attB site. When site specific integration is accomplished, the cell line switches color (from mRuby2 to tagBFP) and resistance (Hygromycin to Blasticidin).

To test the landing pad, we integrated a transcriptional unit that contains H2B fused to mAzamiGreen, driven by the CAG promoter (Fig. 4a). Upon integration of the transfer vector into the hAAVS1 LP, we noted the decrease in mRuby2 expression together with expression of mAzamiGreen and tagBFP in most cells (Fig. 4b), with a small fraction of cells showing no or incorrect integration of the transfer vector. Gratifyingly, clone #2, in which both wild type alleles of hAAVS1 locus were replaced by the landing pad construct, showed two populations as measured by fluorescence upon integration of the transfer vector. This suggested that these two populations had one or two copies that integrated into the genome, further demonstrating the efficiency of integrating into the landing pad (Supplementary Fig. 3b).

While our results have shown the functionality of the landing pad in one genomic locus, this system is by design generalizable to additional loci of interest or additional species by simply replacing the homology arms in the CRISPR-Cas9 vector. It is also straightforward to replace resistance and fluorescent protein markers in both the landing pad and transfer vectors, as these vectors are generated by GG assemblies of parts already existing in the MTK library.

### MTK allows for rapid, one step combinatorial construction of libraries

When designing experiments that require analysis of a large number of protein or circuit variants, such as protein expression or activity optimization and CRISPR/Cas9 screens, it is vital to ensure that the cloning protocol does not present any bottlenecks in terms of throughput, variant representation and diversity. The combinatorial and hierarchical assembly of MTK provides an efficient method to parallelize the construction of large libraries of vectors in one-step combinatorial reactions. Furthermore, it enables such libraries to be reused and delivered repeatedly and in multiple ways.

In order to showcase the ability of MTK to generate libraries of transcriptional unit variants, we performed two combinatorial reactions: First, a library of fluorescent proteins that vary in their localization, and which can be integrated as a single-copy into the BxBI landing pad site; second, a library of viral-delivered sgRNAs that target Cas9 to two fluorescent proteins leading to their disruption.

In the first library, we combined one of three localization tags (NLS (nucleus), NES (cytoplasm) and Lyn (plasma membrane)) with four fluorescent proteins (tagBFP, mAzamiGreen, mScarlet and iRFP713) (Fig. 5a). Performed in parallel, this reaction generated 4 fluorescent variants per localization tag. These libraries of variants were further combined, and pooled together in equimolar amounts with a Landing Pad destination vector. This single reaction created a library of 64 distinct variants, where each vector in the library encodes three random fluorescent proteins that are localized in the nucleus, cytoplasm and plasma membrane. The final library was delivered to the hAAVS1-LP HEK293T cell line described before (Fig. 4) to ensure single-copy integration in every cell. We visualized the expression and localization of the fluorescent proteins after selection with Blasticidin and observed the successful random delivery of no more than 3 fluorescently-tagged subcellular compartments (Fig. 5b), as predicted from the cloning strategy.

**Fig. 5.**
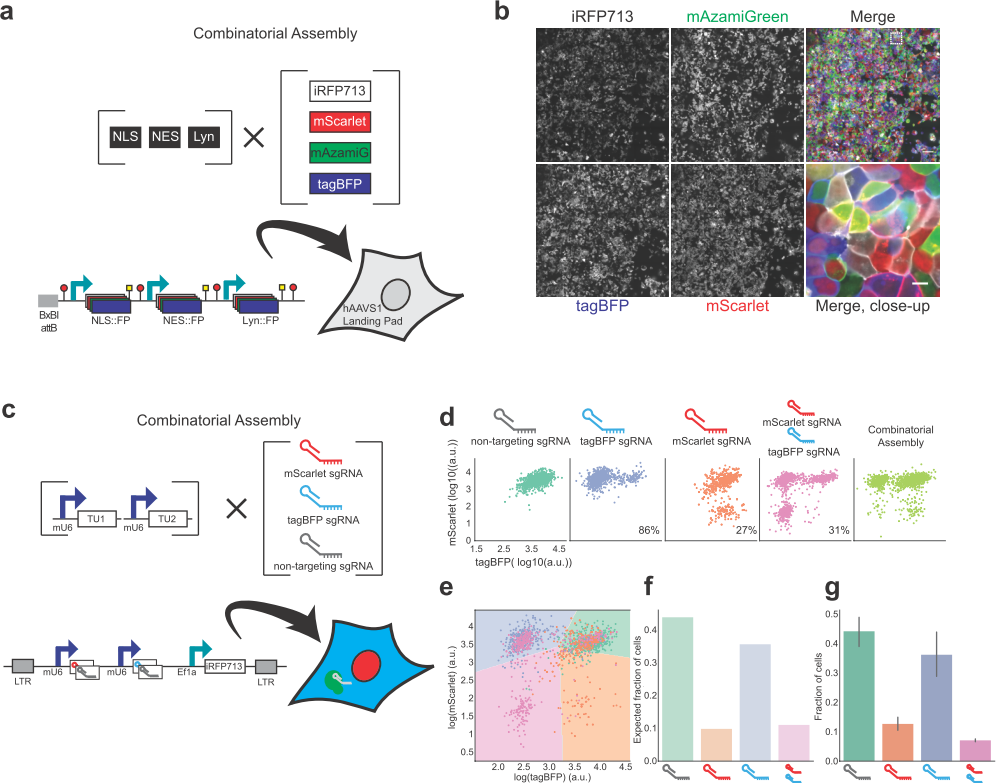
Combinatorial assembly of complex libraries using the MTK. **(a)** Overview of the MTK strategy for a combinatorial, one pot, generation of a library containing 12 combinations of 4 different fluorescent proteins (FP) targeted to three cellular locations. The final multi-TU construct is composed of 3 TUs: EF1a driving the nuclear localization of an FP; CMV expressing the cytoplasmic localization of an FP; and CAG producing a membrane-tethered FP. Each FP is one of 4 variable fluorophores (tagBFP, mAzamiGreen, mScarlet and iRFP713), giving a total of 64 possible variants. Constructs are delivered to the landing pad in HEK293T cells. **(b)** Confocal image of HEK293T+LP, transfected with pooled library in panel (a). iRFP713 shown in white, mScarlet shown in red, mAzamiGreen shown in green and tagBFP shown in blue. Merge and close-up show cells expressing 3 FPs in three subcellular locations displaying the variety of expected combinations. **(c)** Overview of the MTK strategy for a combinatorial, one pot, library of lentiviral vectors carrying sgRNAs targeting tagBFP, mScarlet, both or non-targeting. The final multi-TU construct is composed of three TUs: mU6 promoter driving the expression of mScarlet or non-targeting sgRNA; mU6 promoter driving the expression of tagBFP or non-targeting sgRNA; EF1a driving the expression of iRFP713 for identification of cells that have integrated the construct. Library was produced and transduced to HEK293T cells expressing tagBFP, mScarlet and Cas9 fused to mAzamiGreen (3C cells). **(d)** Scatter plots of tagBFP and mScarlet fluorescence in populations of 3C cells where sgRNAs were individually expressed to target tagBFP, mScarlet, tagBFP and mScarlet, or with non-targeting sgRNA. Last panel shows the scatter plots of tagBFP and mScarlet fluorescence in populations transfected with combinatorially assembled library of sgRNAs. **(e)** Scatter plot of tagBFP and mScarlet fluorescence in populations of 3C cells, where an equal number of cells expresses sgRNAs that target tagBFP, mScarlet, tagBFP and mScarlet, or with non-targeting sgRNA. Shaded areas correspond to 4 classes identified by a linear classifier (green, non-targeting; blue, tagBFP; orange, mScarlet; pink, mScarlet and tagBFP). **(f)** Expected fraction of cells expressing different guide combinations as identified by linear classifier in panel (e) where 4 sgRNA combinations have equal ratios. **(g)** Measured fraction of cells expressing different guide combinations in combinatorial assembly is similar to (f). Bar plot and error bars represents mean and 95% CI of four biological repeats. NLS, Nuclear localization signal; NES, nuclear export signal; Lyn, plasma membrane tag; mAzamiG, mAzamiGreen

Next, we sought to explore whether the MTK facilitates the construction of combinatorial CRISPR knockout libraries. To do that, we generated a library of sgRNAs that target two fluorescent proteins. We first assembled TUs that contain sgRNAs that target tagBFP, mScarlet, or that are non-targeting^17^. We combined those in a multi-TU unit so that a final Lentiviral delivery plasmid contains 2 guide RNAs, ensuring that we had any one of four possible outcomes (tagBFP knockout, mScarlet knockout, tagBFP and mScarlet knockout, or non-targeting control) in the final library (Fig. 5c). Additionally, each lentiviral vector encodes EF1a-driven copy of iRFP713 to facilitate the identification of cells that have been successfully transduced with the sgRNA library.

Since sgRNA targeting has variable efficiency^18^, we concurrently generated four individual lentiviral plasmids for the four outcomes of the library as a control to compare the results of the combinatorial assembly, which would generate all four outcomes in one experiment. We independently transduced the four control viruses in HEK293T cells expressing tagBFP, mScarlet and spCas9 (3C cell line) and quantified tagBFP and mScarlet expression two weeks after transduction. Cells that contained the non-targeting sgRNAs expressed high levels of both tagBFP and mScarlet (Fig. 5d, first panel). Cells expressing the tagBFP-targeting sgRNA showed reduced expression of tagBFP in 86% of the population (Fig. 5d, second panel), while only 27% of cells showed reduced expression of mScarlet when its guide was expressed (Fig. 5d, third panel). Accordingly, when both guides were co-expressed, only 31% cells show reduced expression for both proteins, while most cells had reduced tagBFP (Fig. 5d, fourth panel).

We used these results to build a linear classifier that would allow us to use the observed tagBFP and mScarlet expression to determine the likely sgRNA that each cell received in a population transduced with the full sgRNA library. Due to the inefficient targeting of the mScarlet guide, the linear classifier had an average precision of 0.62 and an average recall of 0.58 (Fig. 5e, Supplementary Fig. 4). This classifier, which was trained on the individual targeted variants of sgRNA, predicted higher than expected proportions of non-targeting and tagBFP sgRNA-expressing cells when presented with an equal proportion of each variant of the library. This is in accordance with the inefficiency of the mScarlet sgRNA targeting (Fig. 5f).

To test the combinatorial assembly approach, we independently transduced the 3C cell line with 4 biological replicates of the sgRNA library. After 14 days of selection, we measured tagBFP and mScarlet expression and identified cells belonging to the four possible outcomes of the viral library, suggesting that all combinatorial possibilities are achievable through this method (Fig. 5g). Importantly, the fraction of each outcome was in accordance with the predictions of the classifier. Taken together, these data argue that the MTK is able to rapidly generate libraries of vectors that express proteins variants or multiple sgRNAs. Additionally, these libraries can be inserted into the genome by transduction or recombination and the diversity of variants, as well as the correct ratio of each variant in the library, is maintained through all cloning steps.

### MTK facilitates optimization of combinatorial gene circuits for synthetic biology applications

The ability to prototype gene circuits in synthetic biology is contingent on rapidly assessing part efficacy, optimizing and implementing changes for the next iteration. The vast combinatorics present in the MTK library present a fruitful opportunity to prototype, test, and iterate on circuit designs in parallel. To explore this idea, we constitutively expressed deactivated *Streptococcus pyogenes* and *Streptococcus aureus* CRISPR Cas9 (dSpCas9 and dSaCas9, respectively) C-terminally fused to either a fluorescent protein (FP), repressor domain (Krüppel associated box, KRAB), or an activator domain (VP64-p65-Rta, VPR)^19–23^. These protein-effector combinations were targeted to two canonical inducible promoters: the GAL4 Upstream Activating Sequence (UAS) or the Tet Responsive Element (TRE) (Fig. 6a). Each gene circuit was assembled adjacent to a constant TU with constitutive EF1a expression of iRFP713, in parallel. In total, twelve different circuits capable of modulating mAzamiGreen expression. We transiently transfected these circuits into HEK293T cells and normalized changes in mAzamiGreen expression by iRFP713.

As expected, in each of the circuit iterations, expression of mAzamiGreen relative to the iRFP713 control is below basal with the repressor domain, or above basal with the activator domain (Fig. 6b). However, the degree of activation and repression varies considerably across each circuit. More specifically, dSpCas9 achieves approximately 3-fold reduction in fluorescence through the KRAB effector and 30-fold induction in both the UAS and TRE promoters. In contrast, the dSaCas9 fused to VPR is an effective activator increasing expression 30-fold, but when fused to a KRAB domain, dSaCas9-mediated repression is comparable to background FP expression levels. The data also suggests that the TRE promoter exhibits higher basal activity due to its overall greater repression, and lower activation when compared to the UAS promoter. This information, in light of other considerations such as protein size or protospacer adjacent motif (PAM) sequence versatility, expedites the rational design and optimization of Cas9-controlled gene expression circuits through the parallelization of prototyping. As a result, this strategy can serve as a useful reference for further customized circuits, and a demonstration that this prototyping can be scaled to easily screen various combinations of genetic circuits.

### MTK streamlines the generation of endogenous viral circuits

Viruses represent a class of naturally occurring genetic circuits that are particularly amenable to construction using the MTK due to their modular genome organization. In particular, the MTK can be relevant to speed the investigation of emerging viral agents by rapidly “booting up” their genes in a mammalian host. A key example corresponds to the filoviruses, a family of highly pathogenic genetically related viruses that includes the Ebola and Marburg viruses^24^. Infection with these agents can cause rapid onset of severe and often fatal hemorrhagic fever in humans and nonhuman primates. First identified in 1976, they remain a significant human pathogen due to their high fatality rate and periodic outbreaks. While global efforts to contain these outbreaks have resulted in fast-tracking of several experimental treatments, there is currently no approved preventive or therapeutic treatment for EBOV infections.

EBOV is encoded by ∼19 kbp negative sense single stranded RNA genome that contains 3’ non-coding regulatory leader (ldr) and 5’ non-coding regulatory trailer (trl) sequences. These sequences flank a large central region that encodes seven distinct proteins (nucleoprotein, NP; 35 kDa phosphoprotein/polymerase co-factor, VP35; 40 kDa matrix protein, VP40; surface glycoprotein, GP; 30 kDa phosphoprotein/polymerase co-factor, VP35; 24 kDa regulatory protein, VP24; and a large RNA dependent RNA polymerase protein, Lpol) (Fig. 7a). In the late 1990’s a 6-plasmid BSL2 ribonucleoprotein (RNP) minigenome replicon system was developed for EBOV to accelerate research on the transcription and genome replication sub-lifecycle of the virus, and aid antiviral discovery efforts^25,26^. This system requires expression of four viral proteins (NP, VP35, VP30, and Lpol), along with aT7 polymerase, and a T7-driven minigenome reporter construct^25^ (Fig. 7a). Transfection of these 6 plasmids into mammalian cells results in expression of NP, VP35, VP30, Lpol viral proteins, and the T7 polymerase. The viral proteins recognize and associate with the T7-transcribed minigenome reporter RNA template (vRNA (-)) then catalyze transcription (mRNA) and replication (aRNA (+)), resulting in measurable reporter gene expression (Fig. 7a). While the 6-plasmid system has been a valuable tool for initial discoveries into EBOV replication and transcription, it is not readily amenable to systematic studies due to its transient nature, inefficiency (all 6 plasmids must be present in the same cell), and variability that limits scaling. More recently, a stable cell line version of this system has been developed via sequential integration of RNP viral proteins^27^. While these cells displayed a highly stable minigenome RNA-protein complex and more robust minigenome activity than the previously reported 6-plasmid systems, generation of this cell line entailed a time-intensive, multi-step approach that is not amenable for every emerging virus or variants of the same virus.

**Fig. 7.**
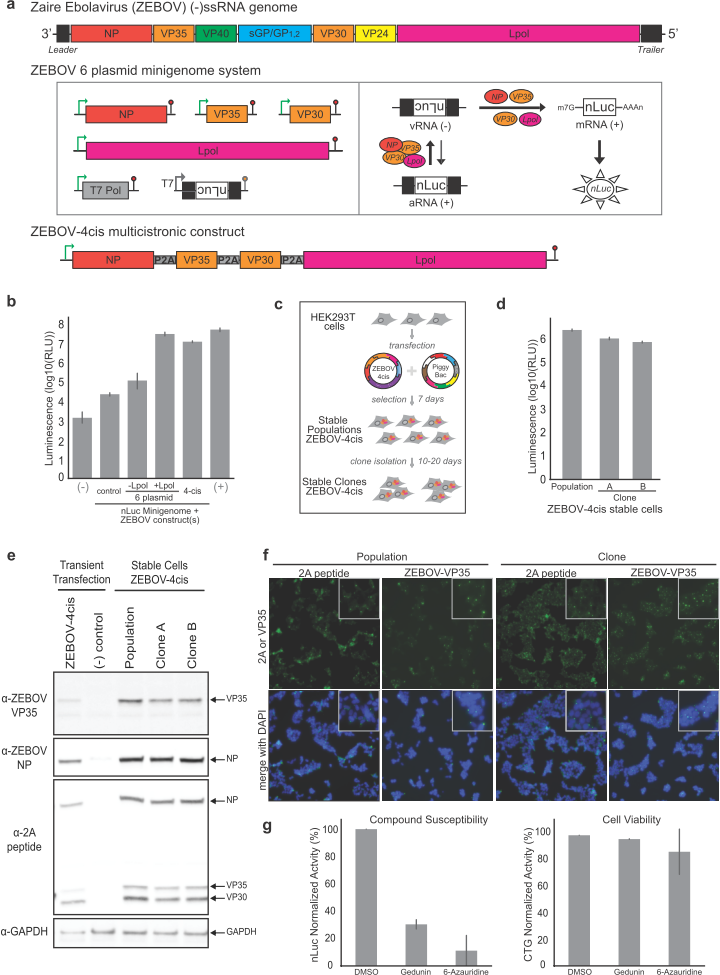
Generating multicistronic constructs for Zaire ebolavirus ribonucleoproteins in a mammalian cell host using the MTK. **(a)** Schematic of the different constructs. Panel 1: Schematic of the Zaire ebolavirus (ZEBOV) negative sense single stranded RNA (-ssRNA) genome. Panel 2: Functional minigenome assay for ZEBOV using the 6 plasmid system that includes expression constructs for NP, VP35, VP30, and Lpol viral proteins as well as the T7 polymerase along with a T7-driven minigenome reporter construct flanked by viral non coding 3’ and 5’ UTRs, the leader and trailer, respectively. Transfection of the 6 plasmids into mammalian cells results in expression of NP, VP35, VP30, Lpol, and T7 and transcription of the minigenome viral RNA (vRNA(-)). The vRNA(-) is replicated to generate antigenomic RNA (aRNA(+)) and transcribed into mRNA by the viral proteins to yield reporter gene expression. Panel 3: Schematic of the Zaire ebolavirus multicistronic construct (ZEBOV-4cis) with NP, VP35, VP30, and Lpol separated by P2A ribosomal skipping site elements. **(b)** Luminescence measurements of ZEBOV minigenome activity. HEK293T cells were transfected with ZEBOV nano luciferase (nLuc) minigenome in combination with pCAGGs empty control plasmid (control), the ZEBOV 6 plasmids (with or without Lpol: +Lpol, -Lpol), or with ZEBOV-4cis in PiggyBac part 0 vector. Positive (+) and negative (-) controls for nLuc expression levels include transfection of only pCAGGS-nLuc plasmid or pCAGGs empty plasmid, respectively. Nano luciferase activity was measured two days post transfection. Bar plots represent the mean of biological replicates (n=2). (**c)** Schematic of ZEBOV-4cis stable cell line generation using PiggyBac transposon. HEK293T cells were co-transfected with MTK043-ZEBOV-4cis and a PiggyBac expression construct for 3 days and selected with hygromycin for 7 days to generate stable cell populations expressing ZEBOV RNP complex proteins. Clones were isolated from these populations via limited dilution plating in additional 10-20 days. (**d**) Luminescence measurements of minigenome activity in ZEBOV-4cis stable population. ZEBOV-4cis stable cells and clones were transfected with a T7-driven ZEBOV minigenome construct encoding an nLuc reporter along with T7 polymerase for 2 days followed by nano luciferase assay. Bar plots represent mean of technical replicates (n=10). **(e)** Western blot confirmation of protein expression. ZEBOV-4cis stable population and clones as well as HEK293T cells transfected with MTK0-43-ZEBOV-4cis were processed for Western blot analysis with mouse anti-ZEBOV VP35, rabbit anti-ZEBOV-NP, mouse anti-2A peptide, and rabbit anti-GAPDH antibodies. **(f)** Immunofluorescence analysis of viral protein localization. ZEBOV-4cis population and clone were stained with with anti-2A peptide and anti-ZEBOV-VP35 primary antibodies and Alexa-488 secondary antibody. DAPI stained nuclei are shown in the merged images. Insets represent 2X magnified fields. **(g)** Effect of chemical compound inhibitors on minigenome activity in ZEBOV-4cis stable cells. ZEBOV-4cis stable population was transfected with a ZEBOV minigenome construct encoding a nano luciferase reporter and treated with DMSO (1%), Gedunin (5uM), or 6-Azauridine (5uM). After 2 days cells were processed for nano luciferase assay for functional minigenome activity and cell titer glo assays for cell viability. Levels of signal relative to DMSO control is plotted. Bar plots represent mean of biological replicates (n=2).

Given the urgent need that outbreaks of emerging viral agents pose to human health, we sought to test if the MTK could be harnessed to streamline and accelerate the generation of BSL2 reagents like the stable EBOV RNP cell lines. We synthesized the EBOV RNP genes and used the MTK multicistronic workflow with Left connector parts that encode P2A ribosomal skipping site elements to rapidly assemble a 4-cistronic construct of Zaire ebolavirus (ZEBOV-4cis) (Fig. 7a) directly and simultaneously into five different Part 0 destination vectors (Supplementary Fig. 5a). These vectors provide distinct types and numbers of genome integration events (via transposase (PiggyBac) or integrases (PhiC31, BxBI)). Digestion followed by gel electrophoresis confirmed high cloning efficiency and correct assembly of ZEBOV-4cis in all 5 destination vectors (Supplementary Fig. 5a). In transient transfections, all ZEBOV-4cis constructs showed no impact on cell viability and displayed levels of minigenome activity similar to the 6-plasmid system and ∼40-60-fold higher activity than the cells lacking Lpol (Fig. 7b, Supplementary Fig. 5b).

We next employed the PiggyBac transposon system to achieve high copy integration of the ZEBOV-4cis construct for a robust functional minigenome reporter readout. HEK293T cells were co-transfected with PiggyBac transposon and the PiggyBac donor construct (MTK043-ZEBOV-4cis) that contains a hygromycin marker cassette, followed by selection with hygromycin for 7 days (Fig. 7c). This resulted in 2 independent stable populations of HEK293T cells expressing ZEBOV RNP proteins, as determined by eGFP minigenome reporter signal (Supplementary Fig. 5c). Clonal cell lines were isolated and expanded from these 2 stable populations in an additional 1-2 weeks. This workflow simplified and speeded the generation of stable EBOV RNP cell lines compared to the previously reported system^27^.

We characterized the resulting EBOV RNP stable cell lines via quantitative minigenome reporter assays, protein expression and localization studies, and analysis of susceptibility to previously described small molecule inhibitors of ZEBOV RNP minigenome activity. ZEBOV-4cis stable populations and clonal cells yielded comparable nLuc minigenome reporter activity (Fig. 7d). ZEBOV viral proteins in both the stable and clonal cell populations were readily detectable with NP and VP35 specific antibodies, and showed expression levels similar to transiently transfected ZEBOV-4cis (Fig. 7e). Additionally, based on the presence of 2A peptide tag that remains at the C-terminus of each of these proteins following “self-cleavage” due to ribosome skipping, NP, VP35, and VP30 were each detectable with an antibody raised against the 2A peptide (Fig. 7e). Larger multi-ORF fusion protein products were not readily detected with the 2A peptide antibodies, indicating efficient cleavage occurred at each of the P2A sites encoded within the ZEBOV-4cis construct. This demonstrates the added benefit that the P2A parts in the MTK provide a potential “universal” method to detect protein expression from multicistronic constructs. This is especially useful for analysis of novel emerging viruses or other proteins where specific antibodies are not available.

Immunofluorescence analysis with VP35 and 2A antibodies show characteristic cytoplasmic punctate structures (Fig. 7f) representing inclusion bodies that are sites of virus replication^28–30^, indicating correct localization of the P2A-tagged viral proteins in these stable cell lines. Due to the lack of available specific antibodies for the Lpol and the lack of a P2A tag in the construct, the Lpol expression cannot be assayed by Western blot or immunofluorescence analysis. However, the fact that these cells support minigenome activity (Fig. 7b and 7d) confirms that Lpol is expressed.

To further characterize the ZEBOV-4cis stable RNP cell line system, we also tested their susceptibility to previously identified EBOV small molecule inhibitors: Gedunin, an inhibitor of heat shock protein 90, and 6-Azauridine, a nucleoside analog^31,32^. ZEBOV-4cis stable cells were transfected with the ZEBOV nLuc minigenome reporter and treated with DMSO (1%, control), Gedunin (5 µM), or 6-Azauridine (5 µM) for 2 days. Luciferase assays revealed that both compounds inhibited minigenome activity by >50%, with minimal impact on cell viability (Fig. 7g). These data indicate that the ZEBOV-4cis stable cell lines generated here are similarly susceptible to known inhibitors identified in transient RNP minigenome systems or recombinant virus systems^31,32^. Therefore, our approach implemented through the MTK introduces a robust workflow for generating BSL2 systems for filoviruses and other viral agents as they emerge, providing a critical opportunity to implement rapid chemical compound susceptibility screens using these reagents.

## Discussion

A key element of cellular engineering for basic science or therapeutic purposes is the ability to quickly build, test, and iterate on designs. This is currently infeasible in mammalian cells using conventional methods in molecular cloning. In this work, we presented the MTK, a platform that takes an important step to remove bottlenecks in mammalian cellular engineering. A major asset of the MTK is a large, characterized suite of modular parts to build versatile TUs. These include promoters, 3’UTRs, insulators, P2A elements and a myriad of coding DNA sequences. We demonstrated how these could be easily combined to construct and recycle TUs of different properties, including different levels of gene expression or multicistronic expression. For rapid implementation of these assembled DNA vectors, we included in the MTK lentiviral, recombinase and Cas9 delivery vectors. These vectors allow the user to choose between transient and stable integration of the same circuits, as well as their copy number. Due to the modularity of the MTK, it is straightforward to generate new variants of these vectors, which we demonstrated by creating BxBI landing pads (LP) for the hAAVS1 locus. These vetted, modular parts constitute fundamental tools for the rapid and facile assembly of genetic circuits.

We illustrated this point in two ways: first, we assembled a combinatorial library of transcriptional units that encode different proteins or guide RNAs; second, we combinatorially assembled new circuits that use different MTK parts. Such libraries are generated through the MTK in a straightforward way, reproducing all variants included in the combinatorial assembly process and maintaining their ratios through all the cloning steps. While our studies tackled a small set of demonstrations, it is clear that many new combinatorial libraries can be built with any parts in the MTK as it currently exists, or as users add to its parts. For example, introducing a panel of C-terminal fluorescent protein tags into the MTK would only require a one-time PCR and sequence verification for each tag to “domesticate” the series as 4a Parts. Additionally, the facile assembly of sgRNAs expedites the production of expression vectors that are easily amenable to CRISPR screens of all scales. With the multitude of ways to multiplex this construction, many TUs and circuits can be built, screened and rapidly prototyped.

Despite the modularity in the MTK, there are situations where caution is needed in using certain parts. For example, on occasion, internal BsaI and BsmBI sites must be removed from coding sequences with a synonymous mutation. For non-coding sequences, it is often difficult to foresee the impact of base switching during domestication of the corresponding part. It is possible to domesticate these parts and retain internal BsaI or BsmBI cut sites with a minor modification of the GG protocol (See “end-on” GG Protocol, Methods). In addition, while the MTK allows flexibility in the assembly of a wide variety of TUs, the sequence of TUs in a final multi-TU plasmid is prescribed at the TU assembly step (e.g. a TU assembled with an LS part 1 will always be the most upstream TU). Therefore, it is prudent to follow general design principles in order to maximize their repurposing^33^, such as assembling TUs with strong constitutive promoters downstream of weaker or inducible promoters to minimize spurious read-through.

The capabilities provided in the MTK are particularly relevant in the context of infectious disease outbreaks where time is of the essence. Recent years have seen outbreaks of new and re-emerging viruses including Middle East respiratory syndrome coronavirus (MERS-CoV), Zika virus, Lassa Fever virus, and Ebola virus, among others, where approved treatments are often lacking. Responding to and containing such outbreaks is an important public health challenge that demands rapid production and iteration of naturally occurring viral circuits in mammalian host cells to enable discovery of inhibitors and investigation of the basic biology of the emerging pathogens. Importantly, given that discovery or availability of a virus sequence does not necessarily correlate with successful culturing of the virus in a lab, the need to test multiple strains and clinical isolates that are also optimized for expression in cell culture is paramount and presents a scaling challenge for conventional cloning approaches. Features unique to the MTK such as multicistronic TU assembly, modular delivery into cells, and facile library generation, in combination with automated workflows, can enable faster iteration over these circuits to quickly identify the optimal reagent for downstream efforts and analyses. Using Zaire Ebolavirus (ZEBOV) as a model, we generated cell lines stably expressing the ZEBOV replication complex components in days, while maintaining a library of these components for future applications and iteration. These cell lines are functional, showing minigenome reporter activity comparable to previously described 6-plasmid transient and stable systems. The ZEBOV-4cis cell lines also exhibit susceptibility to previously identified chemical inhibitors. Thus, the MTK provides a potentially powerful tool for virologists to scale functional experimental studies apace with the recent explosive growth in viral genome sequences^34,35^.

Finally, while not explored in this work, the MTK constitutes a launching platform for additional exciting capabilities. For example, the MTK already expedites the production of sgRNA expression vectors, and with the prevalence of sgRNA screening approaches, we envision incorporating barcoding capabilities that can be read by bulk and single-cell sequencing technologies^36–38^. Furthermore, within the current MTK framework all TUs are read in the same direction. However, it is conceivable to design connector parts (1 and 5) that reverse the orientation of a TU to enable more autonomy in genetic circuit design and optimize transcriptional efficiency^33^. Lastly, the emergence of robotics and high-throughput liquid handling promises an exciting future avenue leveraging automation to streamline assembly. Overall, the MTK represents a significant advancement in cloning infrastructure to further expedite the growth, applications and scope of biological research and biotechnology.

## Acknowledgements

This work was supported by the National Science Foundation Award # 1715108 to H.E-.S and the Defense Advanced Research Projects Agency [grant number HR0011-16-2-0045 to H.E-.S] The content and information does not necessarily reflect the position or the policy of the government, and no official endorsement should be inferred. H.E-.S is an investigator in the Chan Zuckerberg Biohub. This work was also supported by the National Defense Science & Engineering Graduate (NDSEG) Fellowship awarded to A.R.B., A.H.N. and L.C.O. Research performed by G.R.K. and A.L.K. is funded by the Chan Zuckerberg Biohub.

## Author Contributions

J.P.F., A.R.B., and H.E-.S. conceived of the study. A.L.K. and G.R.K. conceived of the virology application. J.P.F., A.R.B., G.R.K., A.H.N., J.T., Q.C.W., E.A., S.Y.C., P.H., and L.C.O. constructed parts library. J.P.F., A.R.B., and G.R.K. collected and processed data. J.P.F., A.R.B., G.R.K., A.L.K., and H.E-.S. interpreted results, wrote and edited the manuscript.

## Methods

### Bacterial Cell Culture

Commercial MachI and XL10 strains (QB3 MacroLab) were used to transform plasmid vectors. A typical transformation mixture consists of 2 μL of the golden gate reaction product, 48 μL bacteria, incubated on ice for 30 minutes, heat shocked at 42°C for 1 minute, recovered on ice for 5 minutes, reaction mixture plated onto selective agar and incubated overnight at 37°C. In the case of multi-TU transformations, cells recovered in LB media for 30 min after heat shock at 37°C before plating reaction onto Kanamycin selective agar plates.

#### Golden Gate Reactions

The general reaction mixture follows 0.5 μL per PCR product, annealed oligos, geneblock, or plasmid (50 fmol.L^-1^); 1 μL T4 DNA Ligase Buffer (10x) (NEB) with 0.25% PEG; 0.5 μL T4 DNA Ligase (NEB) diluted with water to a total volume of 9.5 μL. For BsaI GG reactions, we added 0.5 μL BsaI-HFv2 (NEB #R3733). For BsmBI GG reactions we used 0.5 μL of either BsmBI (NEB) or FastDigest Esp3I (Thermo Scientific FD0454 NEB) (both 10,000 U/mL).

#### Thermocycler protocols

The “GG Long” protocol is primarily used for assembly reactions. The reaction temperature is initially held at 45°C for 2 min to digest the plasmids followed by 20°C for 4 min to anneal constituent parts together. After repeating these first two steps 24 times, the temperature is increased to 60°C for 10 min to digest remaining recognition sites and inactivate the ligase. Then the temperature is held at 80°C for 10 min to inactivate the enzyme. Lastly, the reaction is held at 12°C indefinitely. The “GG End-On” protocol is used when BsaI or BsmBI sites need to be retained in the final product. The temperature is initially held at 45°C for 2 min to digest the plasmid followed by 20°C for 5 min to anneal and ligate the resulting plasmid. These steps are cycled 24 times and then held at 16°C indefinitely.

#### Domestication of Parts

Forward and reverse primers ordered from IDT (www.idtdna.com) were manually designed to anneal to source DNA. Internal BsaI and BsmBI sites were removed and tandem BsaI and BsmBI sites were appended to both the 5’ and 3’ ends of the sequence. PCR was performed with the general reaction mixture of 10 μL Q5 polymerase master mix (NEB MO492S) using 1 μL each of the forward and reverse primers (10 μM) and 0.5 μL of source DNA. The desired PCR product was gel extracted (ThermoFisher) and 1 μL of the final elution added to a BsmBI-mediated GG reaction (described above) with the YTK001 domestication vector. The resulting product was transformed into bacteria and grown in selective LB overnight. Plasmid DNA was extracted (ThermoFisher MiniPrep kit) and sequenced verified.

#### Oligo Annealing

Oligos were designed such that at least 15 bases were complementary and had desired sequence and sticky ends. To anneal oligos into double stranded DNA, we prepared a reaction mixture as follows: 1 μL of each oligo (100 μM), 1 μL T4 Ligase Buffer (NEB), 1 μL T4 PNK, 6 μL water. Mixture was incubated at 37°C for 1 hr and then diluted to a volume of 200 μL. To anneal the oligos, a thermocycler protocol was prepared to hold the temperature of the reaction at 96°C for 6 min, and ramp down 0.1°C per second to 23°C. The reaction is then held at 23°C indefinitely.

#### Geneblocks

Gene fragments were ordered from IDT as either whole constructs or partial constructs with complementary overhangs to ensure proper domestication.

#### TU assembly

Part 1-5 plasmids were pooled together with a recipient Part 678 plasmid following a 2:1 molar ratio in a BsaI-mediated GG reaction as described above.

#### MTU Assembly

Constituent transcriptional unit plasmids were pooled in a 2:1 molar ratio with the destination vector in a BsmBI-mediated GG reaction as described above.

### Mammalian Cell Culture and Transfection

HEK293T and 3T3 cells were maintained in DMEM (Dulbecco’s Modified Eagle Medium, Gibco) supplemented with 10% Fetal Calf Serum (SAFC) and passaged every ∼3 days. Transfections for transient and recombinase-based expression were done in quadruplicate, with Lipofectamine 2000 (Invitrogen), according to manufacturer’s instructions. Transfections for Zaire ebolavirus RNP assays were performed in duplicate or triplicate using standard calcium phosphate transfection methods.

### Lentiviral Production

For lentiviral generation, 24h before transfection, 5E5 HEK293T cells were plated in a 6 well plate containing 2mL of growth media. Equimolar amounts of transfer vector and packaging vectors (pCMV-dR8.91 and pCMV-VSV-G), with a total amount of 4µg DNA, were transfected using Lipofectamine 2000 (Invitrogen), according to manufacturer’s instructions. Media was changed after 16h and virus were collected 48h after transduction, by filtering supernatant through a 0.45µm filter (Millipore SLHV033RS).

For transductions, 1E5 target cells were infected with 1mL, 100µL or 10µL of viral supernatant supplemented with 4ug/mL of polybrene (SCBT sc-134220)

### Flow Cytometry and Data Analysis

For the analysis of promoter, 3’UTR expression and hAAVS1 LP expression, cells were collected in 96 well plates (Corning) and measured using an LSR2 flow cytometer (BD). mAzamiGreen (excitation at 488 nm, emission at 530 nm), mRuby2 or mScarlet (excitation at 561 nm, emission between 610 and 620 nm) and tagBFP (excitation at 355 nm, emission at 450 nm) fluorescence levels were recorded for 10000 events. Gating of single cells, normalization of fluorescence levels, and statistical analysis was performed with custom python scripts.

### Microscopy and Image Processing

For the imaging of combinatorial assembly of the fluorescent protein and localization tags library, 5E4 HEK293T cells were plated in an 8-Well µ-Slide (Ibidi) that contains 200µL of growth media. After 24h and before imaging, growth media was replaced with 200µL of Fluorobrite DMEM (Gibco). Imaging was performed in a temperature and atmosphere controlled chamber on a Zeiss microscope equipped with a Yokagawa CSUX1-A1N-E confocal spinning disk. Images were collected with a 40 x 1.1 NA water immersion objective and Photometrics Evolve 512 EMCCD camera. Images were stitched (ZEN, Zeiss) and gamma-corrected (FIJI) for perception enhancement.

For the imaging of multicistronic fluorescent proteins, 5E4 HEK293T cells were plated in 8-Well µ-Slide (Ibidi) containing 200µL of growth media. Imaging was performed in a temperature and atmosphere controlled chamber on a Nikon Ti Microscope equipped with a Andor Borealis CSU-W1 confocal spinning disk. Images were collected through a 20x 0.75NA air objective, using an Andor 4 Laser Launch for tagBFP (excitation at 405nm, collection between 425 and 475nm), mAzamiGreen (excitation at 488nm, collection between 500 and 550nm), mScarlet (excitation at 561nm, collection between 590 and 650nm) and iRFP713 (excitation at 640nm, collection between 665 and 736nm). An Andor Zyla 4.2 sCMOS was used to detect the images and pixel size was 325nm.

### Viral DNA sequences and Plasmids

Synthetic cDNA sequences for viral NP, VP35, VP30, and Lpol from Zaire ebolavirus Mayinga 1976 isolate (Accession: NC_002549, H.sapiens-tc/COD/1976/Yambuku-Mayinga) were codon optimized (NP, VP35, VP30) and synthesized by IDT, followed by Gibson assembly into the pCAGGs vector backbone. These yielded pCAGGs-NP, pCAGGs-VP35, pCAGGs-VP30, and pCAGGs-Lpol expression plasmids. In order to generate the 4-cistronic (ZEBOV-4cis), viral proteins were PCR amplified from pCAGGs expression vectors and domesticated into MTK0_027 entry vector using the BsmbI site followed by verification with Sanger sequencing. The T7-driven viral minigenome construct p2.0_3E5E_eGFP was a gift from Elke Mühlberger (Addgene plasmid # 69359; http://n2t.net/addgene:69359;RRID_Addgene_69359), which was modified via NdeI and NotI restriction sites to generate p2.0-3E5E-nLuc. T7opt in pCAGGS was a gift from Benhur Lee (Addgene plasmid # 65974; http://n2t.net/addgene:65974;RRID_Addgene_65974).

### Compounds

Gedunin (CAS 2753-30-2) (Fisher Scientific # 33871) and 6-Azauridine (Sigma # A1882-1G) were resuspended in DMSO to generate a 1mM stock solution. Stock solutions were serially diluted in DMSO and media to treat cells with a final concentration of 5µM in 1% DMSO.

### Western blot and Immunofluorescence analyses

Cell lysates were prepared in RIPA lysis and extraction buffer (Thermo #89900) containing protease inhibitors (Sigma #P8340) for Western blotting. Lysates were resolved by SDS-polyacrylamide gel electrophoresis (PAGE), transferred to a polyvinylidene difluoride (PVDF) membrane, and subjected to Western blotting using primary antibodies: rabbit polyclonal Zaire ebolavirus NP (IBT Bioservices #0301-045) (1:1000 dilution), mouse monoclonal Zaire ebolavirus VP35 (Kerafast # EMS702) (1:1000 dilution), mouse monoclonal 2A peptide (Novus Biologicals # NBP2-59627) (1:2000 dilution), Rabbit polyclonal GAPDH (Thermo Fisher Scientific # PA1-987) (1:5000 dilution), and secondary antibodies: goat anti-rabbit or goat anti-mouse polyclonal IRDye-800CW antibodies (VWR, 1:5000 dilution). For immunofluorescence analysis, cells seeded in 12-well tissue culture plates for 24 h were fixed in 4% formaldehyde for 20 min, incubated in permeabilization buffer (1% [vol/vol] Triton X-100 and 0.1% [wt/vol] sodium citrate in PBS) for 10 min and then in blocking buffer (1% [vol/vol] Triton X-100, 0.5% [vol/vol] Tween 20, and 3% bovine serum albumin in PBS) for 30 min. Cells were then incubated overnight with primary antibodies for Zaire ebolavirus VP35 or 2A peptide, secondary antibodies goat anti-mouse or goat anti-rabbit Alexa-488 for 30 min, and stained with DAPI solution (GeneTex # GTX16206) for 10 mins. Cells were imaged at 10X (Leica light microscope).

### Minigenome reporter assays (luciferase, GFP)

ZEBOV minigenome reporter construct (p2.0-3E5E-nLuc, 250 ng) and T7-expression plasmid (pCAGGs-T7opt, 250 ng) were co-transfected with four ZEBOV expression plasmids (pCAGGs-ZEBOV-NP, pCAGGs-ZEBOV-VP35, pCAGGs-ZEBOV-VP30, pCAGGs-ZEBOV-Lpol; 250 ng each) or multicistronic constructs (ZEBOV-4cis, 1000 ng), and pCAGGs empty vector for a total of 1500 ng DNA per 0.5E6 HEK293T cells. Suspension transfected cells were seeded into 8 wells of 96-well plates for 2 days in duplicates. Nano luciferase assays for minigenome function and cell titer glo assays for cell viability were performed as per manufacturer’s instructions (Promega). For minigenome assays in ZEBOV-4cis stable cells, 500 ng each of p2.0-T7-3E5E-nLuc or p2.0-T7-3E5E-eGFP and 500ng of T7-expression plasmid were co-transfected. Nano luciferase levels were assayed as above and eGFP signal was captured via microscopy (Leica, 4X magnification).

