## Supplementary material for "A Toolkit for Rapid Modular Construction of Biological Circuits in Mammalian Cells"

Supplementary Fig. 1. **Comparison of constitutive promoters across cell lines and delivery methods.** **(a)** The mAzamiGreen expression from transient transfection and PiggyBac integration of each promoter was assessed in HEK293T and **(b)** NIH3T3 cells. **(c)** mAzamiGreen expression of each promoter was compared between HEK293T and NIN3T3 in transient expression and **(d)** PiggyBac integration. Each point represents the mean of four biological replicates and error bars represent the standard deviation across replicates.

Supplementary Fig. 2. **Comparison of impact of 3' UTRs across cell lines and delivery methods.** **(a)** The mAzamiGreen expression from transient transfection and PiggyBac integration delivery of each 3' UTR was assessed in HEK293T and **(b)** NIH3T3 cells. **(c)** mAzamiGreen expression of each 3' UTR was compared between HEK293T and NIN3T3 in transient expression and **(d)** PiggyBac integration. Each point represents the mean of four biological replicates and error bars represent the standard deviation across replicates.

Supplementary Fig. 3. **Generation of landing pads for human cell lines.** **(a)** PCR products from landing pad genotyping. P1+P3 indicate presence of WT hAVVS1 locus and P2+P3 indicate presence of BxBI landing pad in hAAVS1 locus. mRuby2, mAzamiGreen and tagBFP expression in populations of parental, Landing pad and Landing Pad with Transfer vector HEK293T cells. **(b)** mRuby2, mAzamiGreen and tagBFP expression in populations of parental, Landing pad #2 and Landing Pad #2 with Transfer vector HEK293T cells. In this clone, both wild type alleles of hAAVS1 locus were replaced by the landing pad construct, showing two populations as measured by fluorescence upon integration of the transfer vector and suggesting these two populations had one or two copies that integrated into the genome. mRuby2 expression indicates presence of hAAVS1 landing pad. mAzamiGreen and tagBFP expression indicates precise integration of transfer vector in hAAVS1 landing pad.

Supplementary Fig. 4. **Building a linear classifier to distinguish target sgRNA knockdown populations. (a)** Classification report of linear classifier. **(b)** Confusion matrix of linear classifier.

Supplementary Fig. 5. **Generation and quality control of multicistronic constructs for Zaire ebolavirus ribonucleoproteins**. **(a)** Multicistronic construct cloning efficiency. ZEBOV multicistronic construct containing 4 viral ORFs separated by P2A elements (ZEBOV-4cis) was BsmBI assembled directly into donor vectors for genome engineering. Specifically, ZEBOV-4cis was generated in donor vectors for PiggyBac transposon (MTK0-43), PhiC31 Integrase (MTK0-13, MTK0-16 tagBFP), and BxBI Integrase (MTK0-14, MTK0-17 tagBFP). The number of bacterial colonies screened and positive for correct construct by size (NotI digestions) is indicated. **(b)** Luminescence measurements of ZEBOV minigenome activity and cell viability. HEK293T cells were transfected with ZEBOV nLuc minigenome in combination with pCAGGs empty control plasmid (control), the ZEBOV 6 plasmids (with or without Lpol: +Lpol, -Lpol), or with ZEBOV-4cis in various part 0 donor vectors. Positive (+) and negative (-) controls include transfection of only pCAGGS-nLuc plasmid or pCAGGs empty plasmid, respectively. Nano luciferase activity was measured two days post transfection. Bar plots represent the mean of biological replicates (n=2). (**c**) GFP analysis of minigenome activity in stable cells. ZEBOV-4cis stable populations and clones were transfected with a T7-driven ZEBOV minigenome construct encoding the eGFP reporter along with T7 polymerase. After 2 days cells were imaged for GFP detection (Leica, 4X).

Figure S1

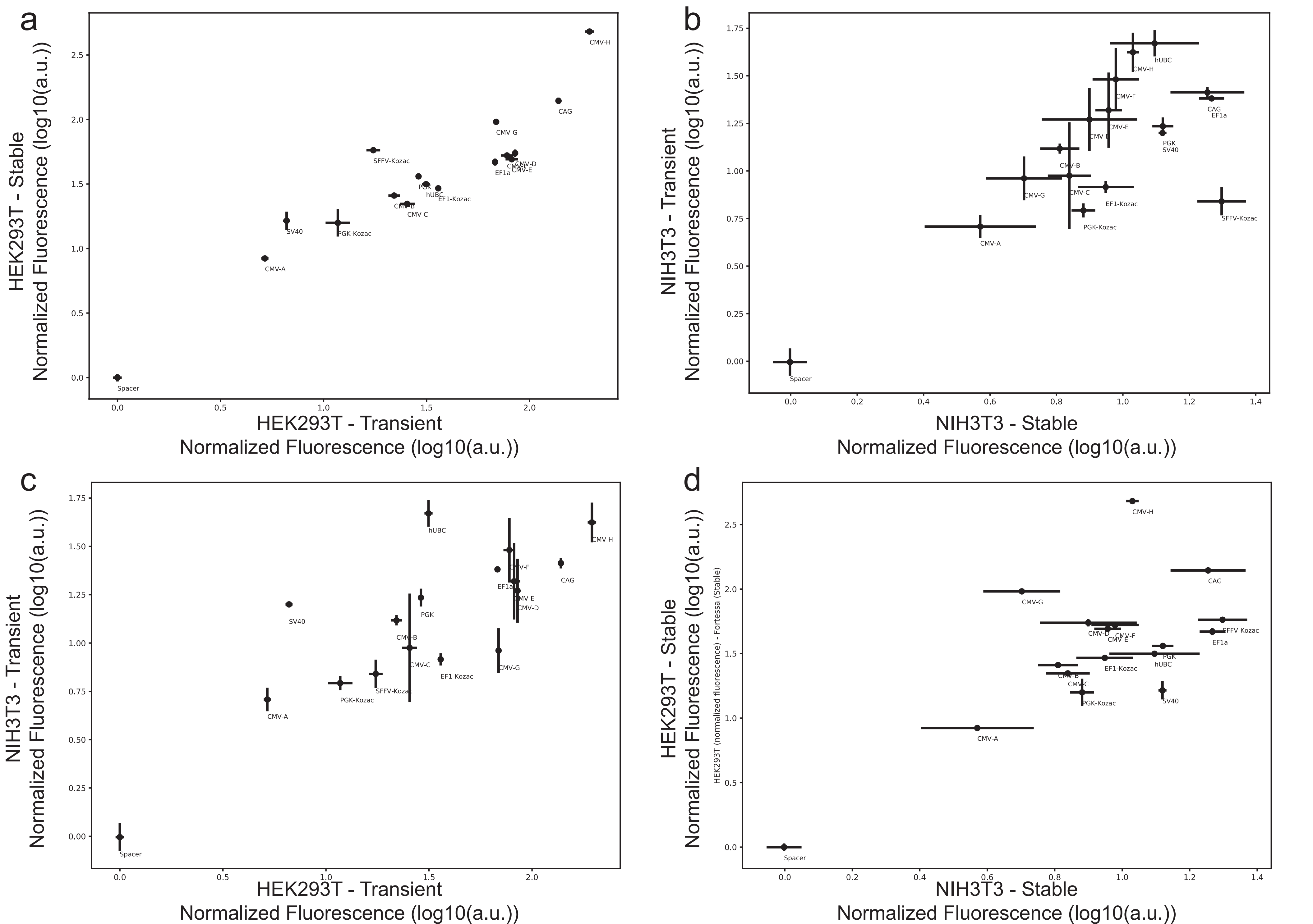
a
HEK293T - Stable
Normalized Fluorescence (log10(a.u.))
HEK293T - Transient
Normalized Fluorescence (log10(a.u.))
Spacer
CMV-A
SV40
PGK-Kozac
SFFV-Kozac
CMV-B
CMV-C
PGK
hUBC
EF1-Kozac
EF1a
CMV-G
CMV-D
CMV-E
CAG
CMV-H
b
NIH3T3 - Transient
Normalized Fluorescence (log10(a.u.))
NIH3T3 - Stable
Normalized Fluorescence (log10(a.u.))
c
NIH3T3 - Transient
Normalized Fluorescence (log10(a.u.))
HEK293T - Transient
Normalized Fluorescence (log10(a.u.))
d
HEK293T - Stable
Normalized Fluorescence (log10(a.u.))
HEK293T (normalized fluorescence) - Fortessa (Stable)
NIH3T3 - Stable
Normalized Fluorescence (log10(a.u.))

Figure S2

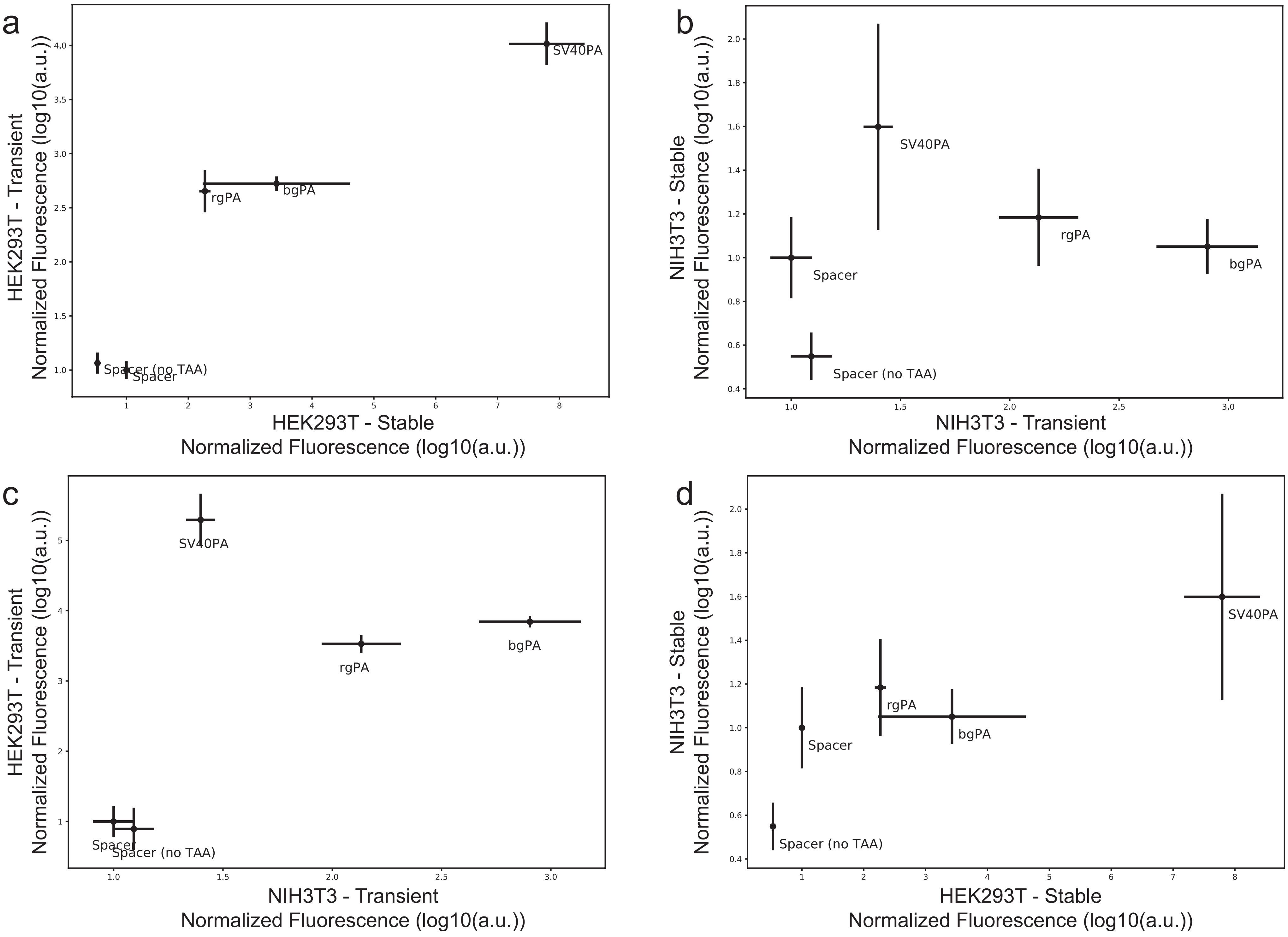
a
HEK293T - Transient
Normalized Fluorescence (log10(a.u.))
HEK293T - Stable
Normalized Fluorescence (log10(a.u.))
SV40PA
rgPA
bgPA
Spacer (no TAA)
Spacer
b
NIH3T3 - Stable
Normalized Fluorescence (log10(a.u.))
NIH3T3 - Transient
Normalized Fluorescence (log10(a.u.))
SV40PA
rgPA
bgPA
Spacer
Spacer (no TAA)
c
HEK293T - Transient
Normalized Fluorescence (log10(a.u.))
NIH3T3 - Transient
Normalized Fluorescence (log10(a.u.))
SV40PA
rgPA
bgPA
Spacer
Spacer (no TAA)
d
NIH3T3 - Stable
Normalized Fluorescence (log10(a.u.))
HEK293T - Stable
Normalized Fluorescence (log10(a.u.))
SV40PA
rgPA
bgPA
Spacer
Spacer (no TAA)

Figure S3

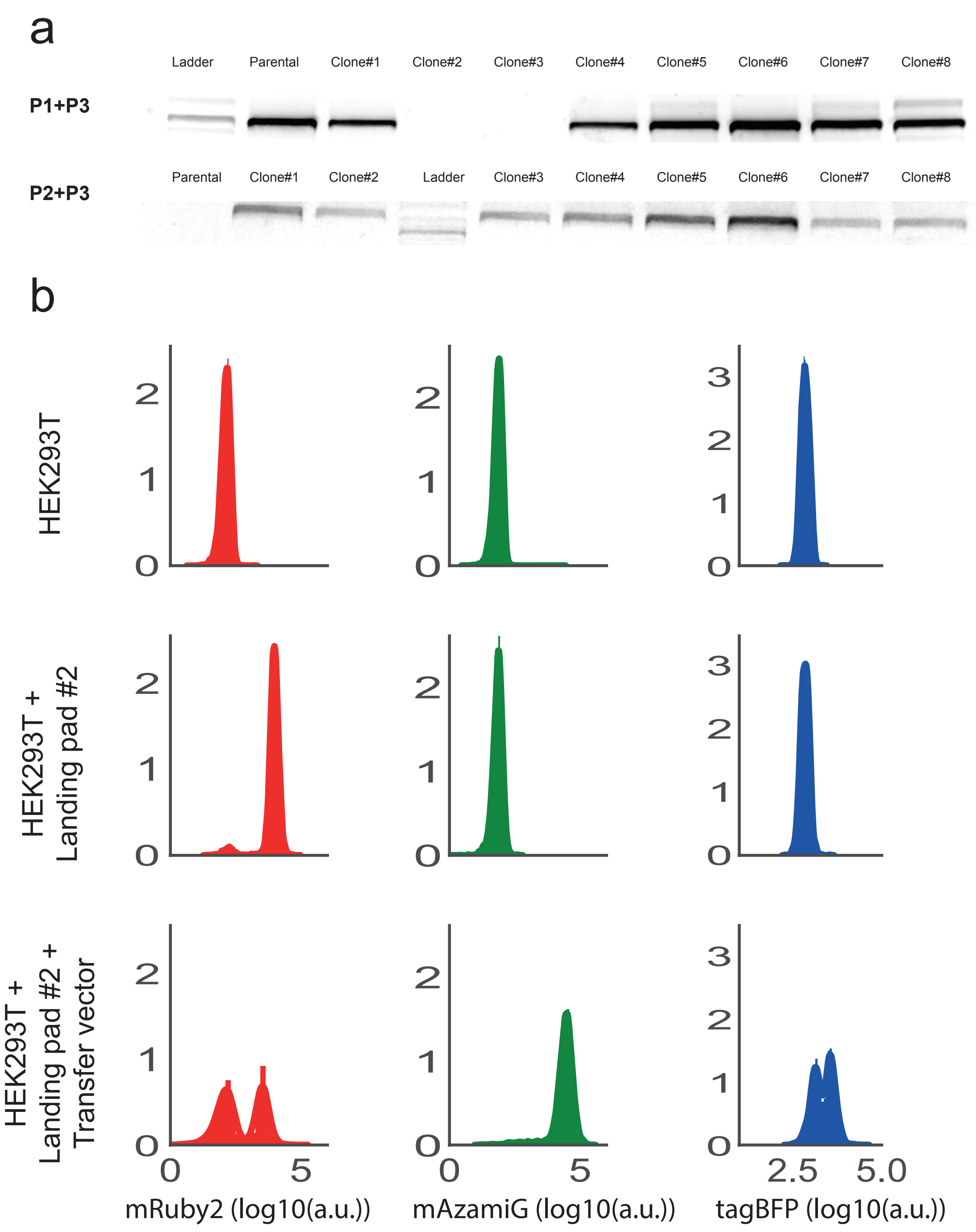
a
Ladder
Parental
Clone#1
Clone#2
Clone#3
Clone#4
Clone#5
Clone#6
Clone#7
Clone#8
P1+P3
Parental
Clone#1
Clone#2
Ladder
Clone#3
Clone#4
Clone#5
Clone#6
Clone#7
Clone#8
P2+P3
b
HEK293T
HEK293T +
Landing pad #2
HEK293T +
Landing pad #2 +
Transfer vector
0
1
2
3
0
5
2.5
5.0
mRuby2 (log10(a.u.))
mAzamiG (log10(a.u.))
tagBFP (log10(a.u.))

Figure S4.

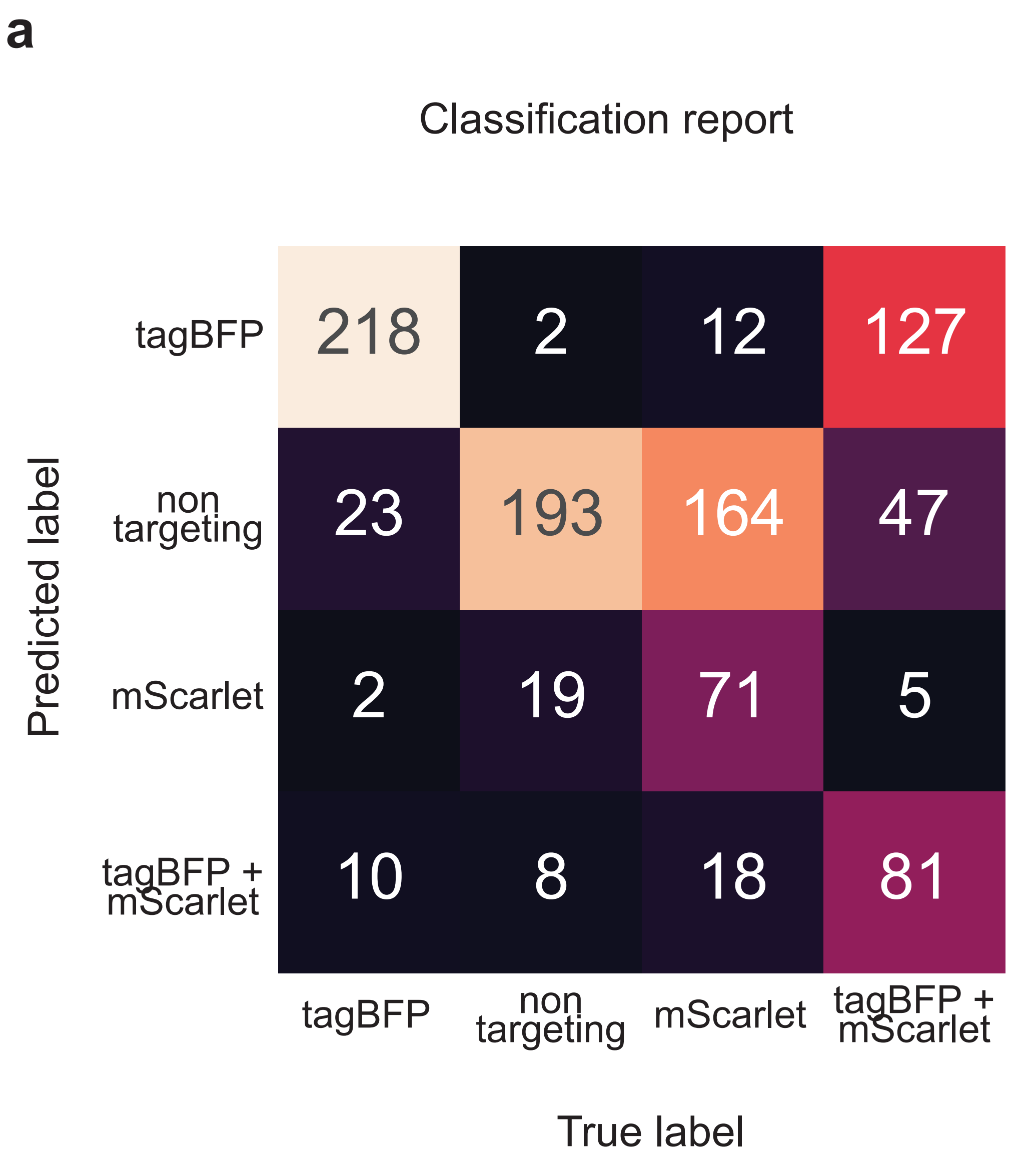
a
Classification report
tagBFP
non targeting
mScarlet
tagBFP + mScarlet
Predicted label
218
2
12
127
23
193
164
47
2
19
71
5
10
8
18
81
tagBFP
non targeting
mScarlet
tagBFP + mScarlet
True label

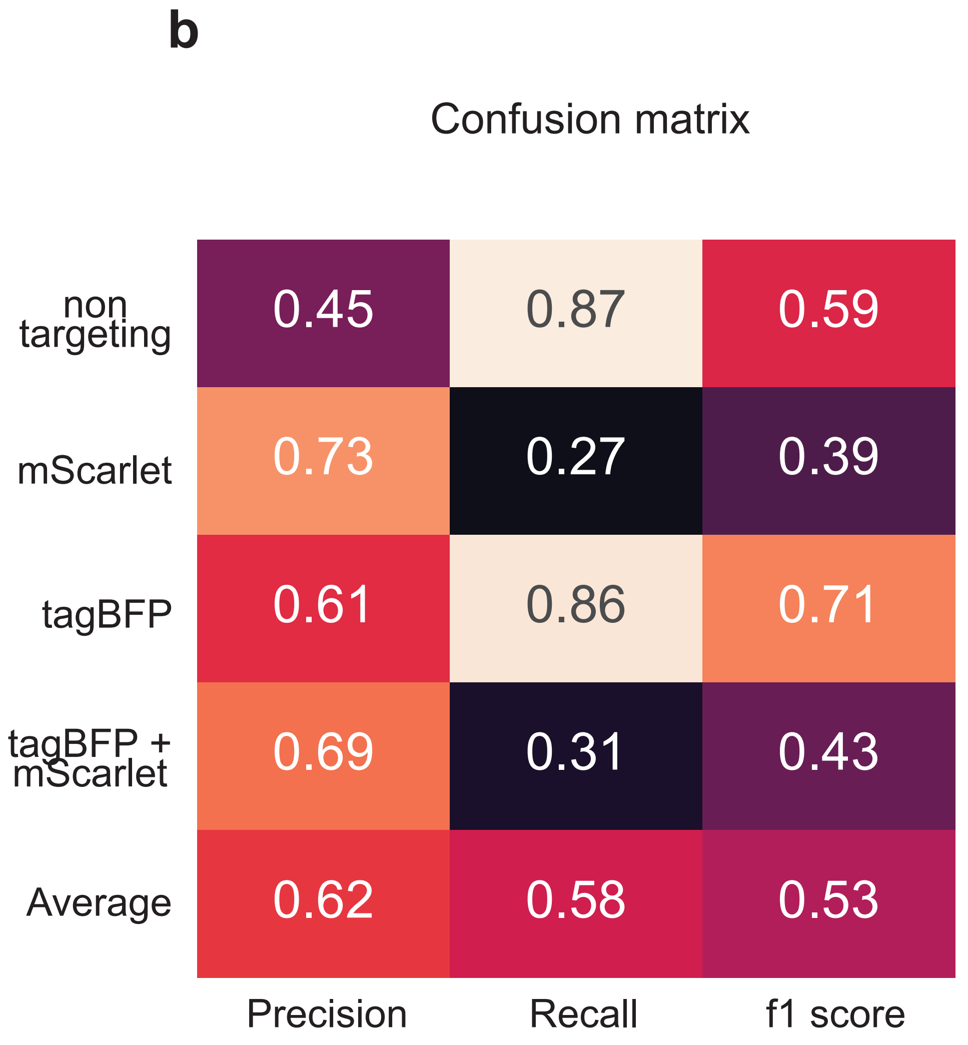
b
Confusion matrix
non targeting
mScarlet
tagBFP
tagBFP + mScarlet
Average
0.45
0.87
0.59
0.73
0.27
0.39
0.61
0.86
0.71
0.69
0.31
0.43
0.62
0.58
0.53
Precision
Recall
f1 score

Figure S5.

**a**

| Donor Vector | Donor Name | ZEBOV-4cis Colonies Screened | ZEBOV-4cis Colonies Correct |
|---|---|---|---|
| MTK0-43 | PiggyBac Transposon | 9 | 9 |
| MTK0-13 | PhiC31 Integrase | 4 | 4 |
| MTK0-14 | BxBI Integrase | 4 | 3 |
| MTK0-16 | PhiC31 tagBFP | 4 | 4 |
| MTK0-17 | BxBI tagBFP | 4 | 4 |

**b**

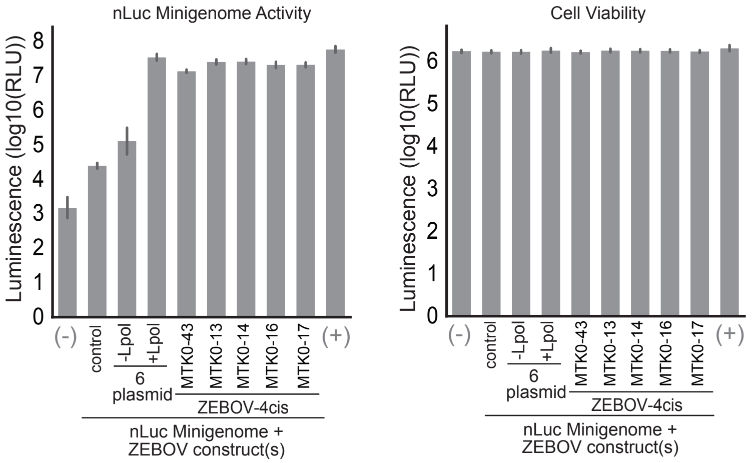


**c**

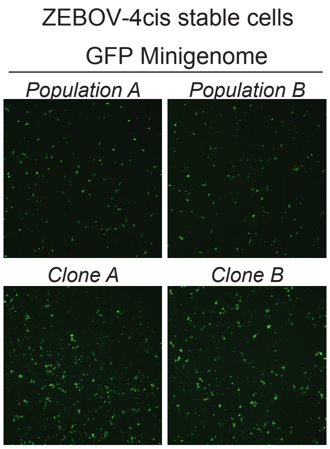

Supplemental Table 1

| Figure | Name | MTK Part Type | Description | Source |
|---|---|---|---|---|
| | MTK1_001 | Part 1 | Connector LS | Lee et al. 2015 |
| | MTK1_002 | Part 1 | Connector L1 | Lee et al. 2015 |
| | MTK1_003 | Part 1 | Connector L2 | Lee et al. 2015 |
| | MTK1_004 | Part 1 | Connector L3 | Lee et al. 2015 |
| | MTK1_005 | Part 1 | Connector L4 | Lee et al. 2015 |
| | MTK1_006 | Part 1 | Connector L5 | Lee et al. 2015 |
| | MTK1_007 | Part 1 | Connector LS' | Lee et al. 2015 |
| | MTK1_008 | Part 1 | Connector LS-2xHS4 | This study |
| | MTK1_009 | Part 1 | Connector L1-2xHS4 | This study |
| | MTK1_010 | Part 1 | Connector L2-2xHS4 | This study |
| | MTK1_011 | Part 1 | Connector L3-2xHS4 | This study |
| | MTK1_012 | Part 1 | Connector L4-2xHS4 | This study |
| | MTK1_013 | Part 1 | Connector L5-2xHS4 | This study |
| | MTK1_014 | Part 1 | Connector LS'-FRT | This study |
| | MTK1_015 | Part 1 | Connector LS-2xHS4-LoxP | This study |
| | MTK1_016 | Part 1 | Connector L6 | Lee et al. 2015 |
| | MTK1_017 | Part 1 | Connector L7 | Lee et al. 2015 |
| | MTK1_018 | Part 1 | Connector L8 | Lee et al. 2015 |
| | MTK1_019 | Part 1 | Connector L6-2xHS4 | This study |
| | MTK1_020 | Part 1 | Connector L7-2xHS4 | This study |
| | MTK1_021 | Part 1 | Connector L8-2xHS4 | This study |
| | MTK1_022 | Part 1 | Connector LS-A4 insulator | This study |
| | MTK1_023 | Part 1 | Connector L1- B1 insulator | This study |
| | MTK1_024 | Part 1 | Connector L2-C3 insulator | This study |
| | MTK1_025 | Part 1 | Connector L3-D1 insulator | This study |
| | MTK1_026 | Part 1 | Connector L4-E1 insulator | This study |
| | MTK1_027 | Part 1 | Connector L5-E4 insulator | This study |
| | MTK1_028 | Part 1 | Connector L6-F1 insulator | This study |
| | MTK1_029 | Part 1 | Connector L7-B2 insulator | This study |
| | MTK1_030 | Part 1 | Connector L1-P2A | This study |
| | MTK1_031 | Part 1 | Connector L2-P2A | This study |
| | MTK1_032 | Part 1 | Connector L3-P2A | This study |
| | MTK1_033 | Part 1 | Connector L4-P2A | This study |
| | MTK1_034 | Part 1 | Connector L5-P2A | This study |
| | MTK2_001 | Part 2 | pTRE | This study |
| | MTK2_004 | Part 2 | pPGK1 | This study |
| | MTK2_005 | Part 2 | pCAG | This study |
| | MTK2_006 | Part 2 | pSV40 | This study |
| | MTK2_007 | Part 2 | pEF1a | This study |
| | MTK2_008 | Part 2 | phUbC | This study |
| | MTK2_012 | Part 2 | pSpacer | This study |
| | MTK2_013 | Part 2 | pSpacer multicistronic | This study |
| | MTK2_014 | Part 2 | pUAS | This study |
| | MTK2_016 | Part 2 | pCMV-A | This study |
| | MTK2_017 | Part 2 | pCMV-B | This study |
| | MTK2_018 | Part 2 | pCMV-C | This study |
| | MTK2_019 | Part 2 | pCMV-D | This study |
| | MTK2_020 | Part 2 | pCMV-E | This study |
| | MTK2_021 | Part 2 | pCMV-F | This study |
| | MTK2_022 | Part 2 | pCMV-G | This study |
| | MTK2_023 | Part 2 | pCMV | This study |
| | MTK2_024 | Part 2 | pPGK1c | This study |

| | | | | |
|---|---|---|---|---|
| | MTK2_025 | Part 2 | pEF1ac | This study |
| | MTK2_026 | Part 2 | pSFFVc | This study |
| | MTK3_002 | Part 3 | Blasticydin Resistance | This study |
| | MTK3_004 | Part 3 | Hygro Resistance | This study |
| | MTK3_005 | Part 3 | Zeomycin Resistance | This study |
| | MTK3_006 | Part 3 | H2B | This study |
| | MTK3_010 | Part 3 | TetOn3G | This study |
| | MTK3_012 | Part 3 | VP16::PYLcs::HA | This study |
| | MTK3_013 | Part 3 | GAL4DBD::ABIcs::Flag | This study |
| | MTK3_022 | Part 3 | mTurquoise2 | Lee et al. 2015 |
| | MTK3_023 | Part 3 | Venus | Lee et al. 2015 |
| | MTK3_024 | Part 3 | mRuby2 | Lee et al. 2015 |
| | MTK3_025 | Part 3 | spCas9 | Lee et al. 2015 |
| | MTK3_027 | Part 3 | VP16::PYLcs-T2A-GAL4DBD::ABIcs | This study |
| | MTK3_031 | Part 3 | p38-mCerulean3 KTR | This study |
| | MTK3_036 | Part 3 | mScarlet | This study |
| | MTK3a_001 | Part 3a | CAXX Lyn | This study |
| | MTK3a_002 | Part 3a | NES | This study |
| | MTK3a_003 | Part 3a | NLS | This study |
| | MTK3a_009 | Part 3a | attP (BxBI) | This study |
| | MTK3a_010 | Part 3a | attP (PhiC31) | This study |
| | MTK3a_011 | Part 3a | EGFP | This study |
| | MTK3a_013 | Part 3a | msfGFP | This study |
| | MTK3a_014 | Part 3a | Krab | This study |
| | MTK3a_015 | Part 3a | VPR-AD | This study |
| | MTK3a_016 | Part 3a | ABI | This study |
| | MTK3a_017 | Part 3a | GAI | This study |
| | MTK3a_018 | Part 3a | H2B | This study |
| | MTK3a_019 | Part 3a | PYL | This study |
| | MTK3a_020 | Part 3a | GID | This study |
| | MTK3a_022 | Part 3a | his::GAI::tagBFP::linker | This study |
| | MTK3a_023 | Part 3a | his::ABI::tagBFP::linker | This study |
| | MTK3a_024 | Part 3a | spCAS9-2xNLS | This study |
| | MTK3a_025 | Part 3a | spdCAS9 (with NLS) | This study |
| | MTK3a_026 | Part 3a | sadCAS9 (with NLS) | This study |
| | MTK3a_027 | Part 3a | tagBFP | This study |
| | MTK3b_003 | Part 3b | T2A | This study |
| | MTK3b_004 | Part 3b | mAzamiGreen | This study |
| | MTK3b_005 | Part 3b | iRFP670 | This study |
| | MTK3b_006 | Part 3b | tagBFP | This study |
| | MTK3b_008 | Part 3b | EBFP2 | This study |
| | MTK3b_009 | Part 3b | mRuby2 | Lee et al. 2015 |
| | MTK3b_010 | Part 3b | mScarlet | This study |
| | MTK3b_011 | Part 3b | iRFP713 | This study |
| | MTK3b_012 | Part 3b | Hygromycin | This study |
| | MTK3b_014 | Part 3b | P2A | This study |
| | MTK3b_018 | Part 3b | SpdCas9 | This study |
| | MTK3b_019 | Part 3b | SadCas9 | This study |
| | MTK3b_021 | Part 3b | VPR AD | This study |
| | MTK3b_022 | Part 3b | KRAB | This study |
| | MTK3b_029 | Part 3b | TET3G | This study |
| | MTK4_001 | Part 4 | Bgh pA | This study |
| | MTK4_002 | Part 4 | SV40 pA | This study |

| | | | | |
|---|---|---|---|---|
| | MTK4_003 | Part 4 | rglb pA | This study |
| | MTK4_004 | Part 4 | Spacer | This study |
| | MTK4_006 | Part 4 | Spacer for multicistronic | This study |
| | MTK4a_001 | Part 4a | CAAX Lyn | This study |
| | MTK4a_002 | Part 4a | NES | This study |
| | MTK4a_003 | Part 4a | NLS | This study |
| | MTK4a_004 | Part 4a | mRuby2 | Lee et al. 2015 |
| | MTK4a_009 | Part 4a | T2A::NLS::EBFP2 | This study |
| | MTK4a_010 | Part 4a | T2A::NLS::mRuby2 | This study |
| | MTK4a_011 | Part 4a | T2A::NLS::iRFP670 | This study |
| | MTK4a_012 | Part 4a | mScarlet | This study |
| | MTK4a_013 | Part 4a | P2A::NLS::iRFP713 | This study |
| | MTK4a_014 | Part 4a | P2A::NLS::mAzamiGreen | This study |
| | MTK4a_015 | Part 4a | msfGFP | This study |
| | MTK4a_016 | Part 4a | P2a-NLS::tagBFP | This study |
| | MTK4a_017 | Part 4a | P2a-NLS::EBFP2 | This study |
| | MTK4a_018 | Part 4a | P2a-NLS::mTurquoise | This study |
| | MTK4a_019 | Part 4a | d2PEST tag | This study |
| | MTK4a_020 | Part 4a | d4PEST tag | This study |
| | MTK4a_021 | Part 4a | T2A::NLS::mRuby2::PEST4d | This study |
| | MTK4a_022 | Part 4a | mTagBFP2 | This study |
| | MTK4a_023 | Part 4a | GAI | This study |
| | MTK4a_024 | Part 4a | ABI | This study |
| | MTK4a_025 | Part 4a | T2A::NLS::mRuby2(noTAA, addGG, for McTU) | This study |
| | MTK4a_026 | Part 4a | HA::2xNLS::ABI | This study |
| | MTK4a_027 | Part 4a | HA::2xNLS::GAI | This study |
| | MTK4b_001 | Part 4b | Bgh pA | This study |
| | MTK4b_002 | Part 4b | SV40 pA | This study |
| | MTK4b_003 | Part 4b | rglb pA | This study |
| | MTK4b_004 | Part 4b | Spacer | This study |
| | MTK4b_006 | Part 4b | Spacer for multicistronic | This study |
| | MTK5_001 | Part 5 | Connector R1 | Lee et al. 2015 |
| | MTK5_002 | Part 5 | Connector R2 | Lee et al. 2015 |
| | MTK5_003 | Part 5 | Connector R3 | Lee et al. 2015 |
| | MTK5_004 | Part 5 | Connector R4 | Lee et al. 2015 |
| | MTK5_005 | Part 5 | Connector R5 | Lee et al. 2015 |
| | MTK5_006 | Part 5 | Connector RE | Lee et al. 2015 |
| | MTK5_007 | Part 5 | Connector RE' | Lee et al. 2015 |
| | MTK5_008 | Part 5 | Connector RE'-2xHS4 | This study |
| | MTK5_009 | Part 5 | Connector R6 | Lee et al. 2015 |
| | MTK5_010 | Part 5 | Connector R7 | Lee et al. 2015 |
| | MTK5_011 | Part 5 | Connector R8 | Lee et al. 2015 |
| | MTK5_012 | Part 5 | Connector RE'-A1 Insulator | This study |
| | MTK6_003 | Part 6 | CMV-Blast-bgPA | This study |
| | MTK6_004 | Part 6 | CMV-tagBFP-T2A-BLAST-bgPA | This study |
| | MTK6_005 | Part 6 | WPRE-Blast | This study |
| | MTK6_006 | Part 6 | Spacer | Lee et al. 2015 |
| | MTK6_007 | Part 6 | attB (phiC31) | This study |
| | MTK6_008 | Part 6 | attB (BxBI) | This study |
| | MTK6_009 | Part 6 | CMV-Hygro-bgPA | This study |
| | MTK6_010 | Part 6 | FRT | This study |
| | MTK6_013 | Part 6 | WPRE-Blast (no terminator) | This study |
| | MTK6_014 | Part 6 | WPRE-tagBFP-T2A-Blast (no terminator) | This study |

| | | | | |
|---|---|---|---|---|
| | MTK6_015 | Part 6 | WPRE | This study |
| | MTK6_016 | Part 6 | WPRE (extended) | This study |
| | MTK7_001 | Part 7 | 3'PB | This study |
| | MTK7_002 | Part 7 | RA_hROSA26 | This study |
| | MTK7_004 | Part 7 | 3' HIV LTR (3rd gen.) | This study |
| | MTK7_005 | Part 7 | 3' AAV ITR | This study |
| | MTK7_006 | Part 7 | FRT | This study |
| | MTK7_007 | Part 7 | Blasticidin - bgPA | This study |
| | MTK7_009 | Part 7 | RA_cROSA | This study |
| | MTK7_010 | Part 7 | BFP-T2A-Blasticidin-BgPa | This study |
| | MTK7_011 | Part 7 | RA_hTha5.10-2-hg18 | This study |
| | MTK7_012 | Part 7 | RA_hCLYBL | This study |
| | MTK7_013 | Part 7 | RA_hAAVS1 | This study |
| | MTK7_014 | Part 7 | 3' HIV LTR (3rd gen.) (extended) | This study |
| | MTK8_004 | Part 8 | F1-AmpR-ColE1 | Lee et al. 2015 |
| | MTK8_005 | Part 8 | F1-KanR-ColE1 | Lee et al. 2015 |
| | MTK8_006 | Part 8 | F1-SpecR-ColE1 | Lee et al. 2015 |
| | MTK8a_001 | Part 8a | AmpR-ColE1 | Lee et al. 2015 |
| | MTK8a_002 | Part 8a | KanR-ColE1 | Lee et al. 2015 |
| | MTK8a_003 | Part 8a | SpecR-ColE1 | Lee et al. 2015 |
| | MTK8a_004 | Part 8a | F1-AmpR-ColE1 | Lee et al. 2015 |
| | MTK8a_005 | Part 8a | F1-KanR-ColE1 | Lee et al. 2015 |
| | MTK8a_006 | Part 8a | F1-SpecR-ColE1 | Lee et al. 2015 |
| | MTK8a_010 | Part 8a | BAC-AmpR | This study |
| | MTK8b_001 | Part 8b | 5'PB | This study |
| | MTK8b_002 | Part 8b | LA_hROSA26 | This study |
| | MTK8b_004 | Part 8b | 5' HIV LTR (3rd gen.) | This study |
| | MTK8b_005 | Part 8b | 5' AAV ITR | This study |
| | MTK8b_006 | Part 8b | LoxP | This study |
| | MTK8b_008 | Part 8b | LA_cROSA | This study |
| | MTK8b_009 | Part 8b | LA_hThal5.10-2-hg18 | This study |
| | MTK8b_010 | Part 8b | LA_hCLYBL | This study |
| | MTK8b_011 | Part 8b | LA_hAAVS1 | This study |
| | MTK234_001 | Part 234 | spacer | Lee et al. 2015 |
| | MTK234_002 | Part 234 | sgRNA-GFPDROPOUT | This study |
| | MTK234_003 | Part 234 | GFP dropout | Lee et al. 2015 |
| | MTK234_005 | Part 234 | sgRNA for cROSA homology arm-1) | This study |
| | MTK234_006 | Part 234 | sgRNA for cROSA Homology arm-2 | This study |
| | MTK234_007 | Part 234 | sgRNA for cROSA Homology arm-3 | This study |
| | MTK234_008 | Part 234 | GFP dropout for multiTU destination vectors | This study |
| | MTK234_010 | Part 234 | TetO-sgRNA | This study |
| | MTK234_012 | Part 234 | hROSA26 sg1 | This study |
| | MTK234_013 | Part 234 | hROSA26 sg2 | This study |
| | MTK234_014 | Part 234 | hROSA26 sg3 | This study |
| | MTK234_015 | Part 234 | hROSA26 sg4 | This study |
| | MTK234_016 | Part 234 | TetO-SasgRNA | This study |
| | MTK234_021 | Part 234 | sgRNA-UAS | This study |
| | MTK234_022 | Part 234 | Thal5.10-2 sgRNA 1 | This study |
| | MTK234_023 | Part 234 | Thal5.10-2 sgRNA 2 | This study |
| | MTK234_024 | Part 234 | Thal5.10-2 sgRNA 3 | This study |
| | MTK234_025 | Part 234 | Thal5.10-2 sgRNA 4 | This study |
| | MTK234_026 | Part 234 | human non targeting sgRNA | This study |
| | MTK234_027 | Part 234 | sgRNA1 -hThal5.10-2_hg18 | This study |

| | | | | |
|---|---|---|---|---|
| | MTK234_028 | Part 234 | sgRNA2 -hThal5.10-2_hg18 | This study |
| | MTK234_029 | Part 234 | sgRNA3 -hThal5.10-2_hg18 | This study |
| | MTK234_030 | Part 234 | sgRNA1 -hCLYBL | This study |
| | MTK234_031 | Part 234 | sgRNA2 -hCLYBL | This study |
| | MTK234_032 | Part 234 | sgRNA1 -hAAVS1 | This study |
| | MTK234_033 | Part 234 | sgRNA2 -hAAVS1 | This study |
| | MTK234_034 | Part 234 | sgRNA3 -hAAVS1 | This study |
| | MTK234_037 | Part 234 | sgRNA-pUAS | This study |
| | MTK234_038 | Part 234 | sgRNA-pSV40 | This study |
| | MTK234_039 | Part 234 | sgRNA1 tagBFP | This study |
| | MTK234_040 | Part 234 | mAzamiGreen sgRNA | This study |
| | MTK234_041 | Part 234 | mScarlet sgRNA | This study |
| | MTK234_043 | Part 234 | sgRNA-pSV40 | This study |
| | MTK234_050 | Part 234 | SaCas9-sgRNA-GFPDROPOUT | This study |
| | MTK234_051 | Part 234 | pUAS sgRNA (Sa) | This study |
| | MTK678_001 | Part 678 | ColE1-AmpR | Lee et al. 2015 |
| | MTK678_002 | Part 678 | BAC-AmpR | This study |
| | MTK0_001 | Part 0 | TU-sgRNA - L1/RE | This study |
| | MTK0_002 | Part 0 | PB Destination - 2xHS4 - HygroR | This study |
| | MTK0_003 | Part 0 | Cas9-sgRNA destination | This study |
| | MTK0_004 | Part 0 | cRosa26 destination | This study |
| | MTK0_005 | Part 0 | hROSA26 Destination - HygroR | This study |
| | MTK0_006 | Part 0 | pLenti Destination vector (3rd generation, low yield) | This study |
| | MTK0_007 | Part 0 | AAV Destination vector | This study |
| | MTK0_008 | Part 0 | hRosa26 landing pad destination vector | This study |
| | MTK0_009 | Part 0 | phiC31 landing | This study |
| | MTK0_010 | Part 0 | BxBI landing | This study |
| | MTK0_011 | Part 0 | hRosa26-PhiC31 landing pad | This study |
| | MTK0_012 | Part 0 | hRosa26-BxBI landing pad | This study |
| | MTK0_013 | Part 0 | Phic31attB destination vector | This study |
| | MTK0_014 | Part 0 | BxBIattB destination vector | This study |
| | MTK0_015 | Part 0 | PB Destination - Insulator + tagBFP | This study |
| | MTK0_016 | Part 0 | Phic31attB tagBFP destination vector | This study |
| | MTK0_017 | Part 0 | BxBI attB tagBFP destination vector | This study |
| | MTK0_018 | Part 0 | TULSR1 - spacer | This study |
| | MTK0_019 | Part 0 | TUL1R2 spacer | This study |
| | MTK0_020 | Part 0 | TUL2R3 spacer | This study |
| | MTK0_021 | Part 0 | TUL3R4 spacer | This study |
| | MTK0_022 | Part 0 | TUL4R5 spacer | This study |
| | MTK0_023 | Part 0 | TUL5R6 spacer | This study |
| | MTK0_024 | Part 0 | TUL6R7 spacer | This study |
| | MTK0_025 | Part 0 | TUL7R8 spacer | This study |
| | MTK0_026 | Part 0 | TUL8RE spacer | This study |
| | MTK0_027 | Part 0 | Part Entry Vector) | This study |
| | MTK0_028 | Part 0 | cRosa26 landing pad destination vector | This study |
| | MTK0_029 | Part 0 | hThal5.10-2 landing pad destination vector | This study |
| | MTK0_030 | Part 0 | phiC31-hygro-mRuby2PEST landing | This study |
| | MTK0_031 | Part 0 | BxBI-hygro-mRuby2PEST landing | This study |
| | MTK0_032 | Part 0 | hThal5.10-2 Destination - HygroR | This study |
| | MTK0_033 | Part 0 | cROSA26 PhiC31 LP | This study |
| | MTK0_034 | Part 0 | hROSA26 PhiC31 LP | This study |
| | MTK0_035 | Part 0 | hThal5.10-2 PhiC31 LP | This study |
| | MTK0_036 | Part 0 | cROSA26 BxBI LP | This study |

| | | | | |
|---|---|---|---|---|
| | MTK0_037 | Part 0 | hROSA26 BxBI LP | This study |
| | MTK0_038 | Part 0 | hThal5.10-2 BxBI LP | This study |
| | MTK0_043 | Part 0 | PB Destination - A1 Insulator - HygroR | This study |
| | MTK0_044 | Part 0 | pLenti Destination vector (w/o insulator) | This study |
| | MTK0_045 | Part 0 | cROSA26 DEST (hygro) | This study |
| | MTK0_046 | Part 0 | hThal5.10-2_hg18 CAS9 Destination - HygroR | This study |
| | MTK0_047 | Part 0 | hCLYBL CAS9 Destination - HygroR | This study |
| | MTK0_048 | Part 0 | hAAVS1 CAS9 Destination - HygroR | This study |
| | MTK0_049 | Part 0 | hThal5.10-2_hg18 landing pad destination vector | This study |
| | MTK0_050 | Part 0 | hCLYBL landing pad destination vector | This study |
| | MTK0_051 | Part 0 | hAAVS1 landing pad destination vector | This study |
| | MTK0_052 | Part 0 | PhiC31 LP on hThal5.10-2-hg18 (Hygro::mRuby) | This study |
| | MTK0_053 | Part 0 | BxBI LP on hThal5.10-2-hg18 (Hygro::mRuby) | This study |
| | MTK0_054 | Part 0 | PhiC31 LP on hCLYBL (Hygro::mRuby) | This study |
| | MTK0_055 | Part 0 | BxBI LP on hCLYBL (Hygro::mRuby) | This study |
| | MTK0_056 | Part 0 | PhiC31 LP on hAAVS1 (Hygro::mRuby) | This study |
| | MTK0_057 | Part 0 | BxBI LP on hAAVS1 (Hygro::mRuby) | This study |
| | MTK0_062 | Part 0 | pHR DEST | This study |
| 2a | JPF0507 | | PB_pPGK1-NLS-mAzamiGreen-bghpA_pCAG-H2B::mScarlet-rglpA | This study |
| 2a | JPF0508 | | PB_pCAG-NLS-mAzamiGreen-bghpA_pCAG-H2B::mScarlet-rglpA | This study |
| 2a | JPF0509 | | PB_pSV40-NLS-mAzamiGreen-bghpA_pCAG-H2B::mScarlet-rglpA | This study |
| 2a | JPF0510 | | PB_pEF1a-NLS-mAzamiGreen-bghpA_pCAG-H2B::mScarlet-rglpA | This study |
| 2a | JPF0511 | | PB_phubc-NLS-mAzamiGreen-bghpA_pCAG-H2B::mScarlet-rglpA | This study |
| 2a | JPF0512 | | PB_spacer-NLS-mAzamiGreen-bghpA_pCAG-H2B::mScarlet-rglpA | This study |
| 2a | JPF0513 | | PB_pCMV-A-NLS-mAzamiGreen-bghpA_pCAG-H2B::mScarlet-rglpA | This study |
| 2a | JPF0514 | | PB_pCMV-B-NLS-mAzamiGreen-bghpA_pCAG-H2B::mScarlet-rglpA | This study |
| 2a | JPF0515 | | PB_pCMV-C-NLS-mAzamiGreen-bghpA_pCAG-H2B::mScarlet-rglpA | This study |
| 2a | JPF0516 | | PB_pCMV-D-NLS-mAzamiGreen-bghpA_pCAG-H2B::mScarlet-rglpA | This study |
| 2a | JPF0517 | | PB_pCMV-E-NLS-mAzamiGreen-bghpA_pCAG-H2B::mScarlet-rglpA | This study |
| 2a | JPF0518 | | PB_pCMV-F-NLS-mAzamiGreen-bghpA_pCAG-H2B::mScarlet-rglpA | This study |
| 2a | JPF0519 | | PB_pCMV-G-NLS-mAzamiGreen-bghpA_pCAG-H2B::mScarlet-rglpA | This study |
| 2a | JPF0520 | | PB_pCMV-NLS-mAzamiGreen-bghpA_pCAG-H2B::mScarlet-rglpA | This study |
| 2a | JPF0521 | | PB_pPGK1c-NLS-mAzamiGreen-bghpA_pCAG-H2B::mScarlet-rglpA | This study |
| 2a | JPF0522 | | PB_pEF1ac-NLS-mAzamiGreen-bghpA_pCAG-H2B::mScarlet-rglpA | This study |
| 2a | JPF0523 | | PB_pSFFVc-NLS-mAzamiGreen-bghpA_pCAG-H2B::mScarlet-rglpA | This study |
| 2b | ARB287a | | PB_pEF1a-NLS-mAzamiGreen-bghpA_pCAG-H2B::mScarlet-rglpA | This study |
| 2b | ARB287b | | PB_pEF1a-NLS-mAzamiGreen-SV40pA_pCAG-H2B::mScarlet-rglpA | This study |
| 2b | ARB287c | | PB_pEF1a-NLS-mAzamiGreen-rglpA_pCAG-H2B::mScarlet-rglpA | This study |
| 2b | ARB287d | | PB_pEF1a-NLS-mAzamiGreen-spacer_pCAG-H2B::mScarlet-rglpA | This study |
| 2b | ARB287e | | PB_pEF1a-NLS-mAzamiGreen-spacermulticistronic_pCAG-H2B::mScarlet-rglpA | This study |
| 3 | JPF0419d | | PB_pCAG-Lyn-iRFP713-P2A-NES-mAzamiGreen-P2A-p38KTR-mCerulean-P2A-H2B::mScarlet | This study |
| 3 | JPF0419v | | PB_pCAG-Lyn-iRFP713_pCAG-NES-mAzamiGreen_pCAG-p38KTR-mCerulean_pCAG-H2B::mScarlet | This study |
| 4 | JPF0446 | | BXBI_pCAG-H2B::msfGFP-bghpA | This study |
| 5a,b | ARB340 | | LS-A4_EF1a-NLS::iRFP713-bgPA_R1 | This study |
| 5a,b | ARB341 | | LS-A4_EF1a-NLS::tagBFP-bgPA_R1 | This study |
| 5a,b | ARB342 | | LS-A4_EF1a-NLS::mAzamiGreen-bgPA_R1 | This study |
| 5a,b | ARB343 | | LS-A4_EF1a-NLS::mRuby2-bgPA_R1 | This study |
| 5a,b | ARB344 | | L1-B1_CMV-NES::iRFP713-rglPA_R2 | This study |
| 5a,b | ARB345 | | L1-B1_CMV-NES::tagBFP-rglPA_R2 | This study |
| 5a,b | ARB346 | | L1-B1_CMV-NES::mAzamiGreen-rglPA_R2 | This study |
| 5a,b | ARB347 | | L1-B1_CMV-NES::mRuby2-rglPA_R2 | This study |
| 5a,b | ARB348 | | L2-C3_CAG-Lyn::iRFP713-SV40_RE | This study |
| 5a,b | ARB349 | | L2-C3_CAG-Lyn::tagBFP-SV40_RE | This study |

| | | | | |
|---|---|---|---|---|
| 5a,b | ARB350 | | L2-C3_CAG-Lyn::mAzamiGreen-SV40_RE | This study |
| 5a,b | ARB351 | | L2-C3_CAG-Lyn::mRuby2-SV40_RE | This study |
| 5c,d | JPF0464 | | pHR_pEF1a-iRFP713_non targeting sgRNA | This study |
| 5c,d | JPF0465 | | pHR_pEF1a-iRFP713_mScarlet sgRNA | This study |
| 5c,d | JPF0466 | | pHR_pEF1a-iRFP713_tagBFP sgRNA | This study |
| 5c,d | JPF0468 | | pHR_pEF1a-iRFP713_mScarletsgRNA_tagBFP sgRNA | This study |
| 5c,d | JPF0454 | | pHR_pPGK1-spCAS9-P2A-mAzamiGreen | This study |
| 5c,d | JPF0455 | | pHR_pPGK1-NES-tagBFP | This study |
| 5c,d | JPF0456 | | pHR_pPGK1-mScarlet | This study |
| 6 | JPF0535a | | PB_pEF1a-tet3G-P2A-iRFP713_pEF1a-spdCAS9::tagBFP-P2A-tagBFP-SV40PA_mu6-sgTRE_pTRE-NLS::mAzamiGreen-rgPA | This study |
| 6 | JPF0535b | | PB_pEF1a-tet3G-P2A-iRFP713_pEF1a-spdCAS9::KRAB-P2A-tagBFP-SV40PA_mu6-sgTRE_pTRE-NLS::mAzamiGreen-rgPA | This study |
| 6 | JPF0535c | | PB_pEF1a-tet3G-P2A-iRFP713_pEF1a-spdCAS9::VPR-P2A-tagBFP-SV40PA_mu6-sgTRE_pTRE-NLS::mAzamiGreen-rgPA | This study |
| 6 | JPF0536a | | PB_pEF1a-tet3G-P2A-iRFP713_pEF1a-spdCAS9::tagBFP-P2A-tagBFP-SV40PA_mu6-sgUAS_pUAS-NLS::mAzamiGreen-rgPA | This study |
| 6 | JPF0536b | | PB_pEF1a-tet3G-P2A-iRFP713_pEF1a-spdCAS9::KRAB-P2A-tagBFP-SV40PA_mu6-sgUAS_pUAS-NLS::mAzamiGreen-rgPA | This study |
| 6 | JPF0536c | | PB_pEF1a-tet3G-P2A-iRFP713_pEF1a-spdCAS9::VPR-P2A-tagBFP-SV40PA_mu6-sgUAS_pUAS-NLS::mAzamiGreen-rgPA | This study |
| 6 | ARB365 | | PB_pEF1a-tet3G-P2A-iRFP713_pEF1a-sadCAS9::mRuby2-P2A-mRuby2-SV40PA_mu6-sgUAS_pUAS-NLS::mAzamiGreen-rgPA | This study |
| 6 | ARB366 | | PB_pEF1a-tet3G-P2A-iRFP713_pEF1a-sadCAS9::KRAB-P2A-mRuby2-SV40PA_mu6-sgUAS_pUAS-NLS::mAzamiGreen-rgPA | This study |
| 6 | ARB367 | | PB_pEF1a-tet3G-P2A-iRFP713_pEF1a-sadCAS9::VPR-P2A-mRuby2-SV40PA_mu6-sgUAS_pUAS-NLS::mAzamiGreen-rgPA | This study |
| 6 | ARB385 | | PB_pEF1a-tet3G-P2A-iRFP713_pEF1a-sadCAS9::tagBFP-P2A-mRuby2-SV40PA_mu6-sgTRE_pTRE-NLS::mAzamiGreen-rgPA | This study |
| 6 | ARB386 | | PB_pEF1a-tet3G-P2A-iRFP713_pEF1a-sadCAS9::KRAB-P2A-mRuby2-SV40PA_mu6-sgTRE_pTRE-NLS::mAzamiGreen-rgPA | This study |
| 6 | ARB387 | | PB_pEF1a-tet3G-P2A-iRFP713_pEF1a-sadCAS9::VPR-P2A-mRuby2-SV40PA_mu6-sgTRE_pTRE-NLS::mAzamiGreen-rgPA | This study |
| 7 | MTK0_013-ZEBOV-4cis | | ZEBOV NP-P2A-VP35-P2A-VP30-P2A-Lpol in MTK0_013 PhiC31attB destination vector | This study |
| 7 | MTK0_014-ZEBOV-4cis | | ZEBOV NP-P2A-VP35-P2A-VP30-P2A-Lpol in MTK0_013 BxBIattB destination vector | This study |
| 7 | MTK0_016-ZEBOV-4cis | | ZEBOV NP-P2A-VP35-P2A-VP30-P2A-Lpol in MTK0_013 PhiC31attB tagBFP destination vector | This study |
| 7 | MTK0_017-ZEBOV-4cis | | ZEBOV NP-P2A-VP35-P2A-VP30-P2A-Lpol in MTK0_013 BxBIattB tagBFP destination vector | This study |
| 7 | MTK0_043-ZEBOV-4cis | | ZEBOV NP-P2A-VP35-P2A-VP30-P2A-Lpol in MTK0_043 PB Destination - A1 Insulator - HygroR | This study |
| 7 | ZEBOV-NP | Part 3 | ZEBOV nucleocapsid | This study |
| 7 | ZEBOV-VP35 | Part 3 | ZEBOV viral protein 35 | This study |
| 7 | ZEBOV-VP30 | Part 3 | ZEBOV viral protein 30 | This study |
| 7 | ZEBOV-Lpol | Part 3 | ZEBOV large RNA dependent RNA polymerase | This study |

### Supplemental Table 2

| Figure | Target Locus | Protospacer Sequence | Cas9 species (Part Plasmid) | PAM Sequence | Forward Oligo | Reverse Oligo | Backbone Vector | Source |
|---|---|---|---|---|---|---|---|---|
| 4 | hAAVS1_1 | GAGCCACATTAACCGGCCCT | S. pyogenes (MTK3_025) | GGG | TGTTTGGAGCCACATTAACCGGCCCTG | TAAACAGGGCCGGTTAATGTGGCTCCA | MTK234_002 | This study |
| 4 | hAAVS1_2 | ATTCCCAGGGCCGGTTAATG | S. pyogenes (MTK3_025) | TGG | TGTTTGATTCCCAGGGCCGGTTAATGG | TAAACCATTAACCGGCCCTGGGAATCA | MTK234_002 | This study |
| 4 | hAAVS1_3 | GGGGCCACTAGGGACAGGAT | S. pyogenes (MTK3_025) | TGG | TGTTTGGGGGCCACTAGGGACAGGATG | TAAACATCCTGTCCCTAGTGGCCCCCA | MTK234_002 | This study |
| 5 | tagBFP | CTACAACGTCAAGATCAGAG | S. pyogenes (MTK3_025) | GGG | TGTTTGCTACAACGTCAAGATCAGAGG | TAAACCTCTGATCTTGACGTTGTAGCA | MTK234_002 | This study |
| 5 | mScarlet | CCACAACGAAGATTATACCG | S. pyogenes (MTK3_025) | TGG | TGTTTGCCACAACGAAGATTATACCGG | TAAACCGGTATAATCTTCGTTGTGGCA | MTK234_002 | This study |
| 5 | human non-targeting | ACGGAGGCTAAGCGTCGCAA | S. pyogenes (MTK3_025) | | TGTTTGACGGAGGCTAAGCGTCGCAAG | TAAACTTGCGACGCTTAGCCTCCGTCA | MTK234_002 | This study |
| 6 | TRE | TACGTTCTCTATCACTGATA | S. pyogenes (MTK3b_018) | GGG | TGTTTGTACGTTCTCTATCACTGATAG | TAAACTATCAGTGATAGAGAACGTACA | MTK234_002 | This study |
| 6 | TRE | GTTACTCCCTATCAGTGATA | S. aureus (MTK3b_019) | AGGAGT | TGTTTGGTTACTCCCTATCAGTGATAG | ATAACTATCACTGATAGGGAGTAACCA | MTK234_050 | This study |
| 6 | UAS | GAGCACTGTCCTCCGAACGT | S. pyogenes (MTK3b_018) | CGG | TGTTTGGAGCACTGTCCTCCGAACGTG | TAAACACGTTCGGAGGACAGTGCTCCA | MTK234_002 | This study |
| 6 | UAS | GAACGTCGGAGCACTGTCCT | S. aureus (MTK3b_019) | CCGAAC | TGTTTGGAACGTCGGAGCACTGTCCTG | ATAACAGGACAGTGCTCCGACGTTCCA | MTK234_050 | This study |
